# *ATRAID* regulates the action of nitrogen-containing bisphosphonates on bone

**DOI:** 10.1101/338350

**Authors:** Lauren E. Surface, Damon T. Burrow, Jinmei Li, Jiwoong Park, Sandeep Kumar, Cheng Lyu, Niki Song, Zhou Yu, Abbhirami Rajagopal, Yangjin Bae, Brendan H. Lee, Steven Mumm, Charles C. Gu, Jonathan C. Baker, Mahshid Mohseni, Melissa Sum, Margaret Huskey, Shenghui Duan, Vinieth N. Bijanki, Roberto Civitelli, Michael J. Gardner, Chris M. McAndrew, William M. Ricci, Christina A. Gurnett, Kathryn Diemer, Fei Wan, Christina L. Costantino, Kristen M. Shannon, Noopur Raje, Thomas B. Dodson, Daniel A. Haber, Jan E. Carette, Malini Varadarajan, Thijn R. Brummelkamp, Kivanc Birsoy, David M. Sabatini, Gabe Haller, Timothy R. Peterson

## Abstract

Nitrogen-containing bisphosphonates (N-BPs), such as alendronate, are the most widely prescribed medications for diseases involving bone, with nearly 200 million prescriptions written annually. Recently, widespread use of N-BPs has been challenged due to the risk of rare but traumatic side effects such as atypical femoral fracture (AFFs) and osteonecrosis of the jaw (ONJ). N-BPs bind to and inhibit farnesyl diphosphate synthase (FDPS), resulting in defects in protein prenylation. Yet it remains poorly understood what other cellular factors might allow N-BPs to exert their pharmacological effects. Here, we performed genome-wide studies in cells and patients to identify the poorly characterized gene, *ATRAID*. Loss of *ATRAID* function results in selective resistance to N-BP-mediated loss of cell viability and the prevention of alendronate-mediated inhibition of prenylation. *ATRAID* is required for alendronate inhibition of osteoclast function, and *ATRAID*-deficient mice have impaired therapeutic responses to alendronate in both postmenopausal and senile (old age) osteoporosis models. Lastly, we performed exome sequencing on patients taking N-BPs that suffered ONJ or an AFF. *ATRAID* is one of three genes that contain rare non-synonymous coding variants in patients with ONJ or AFF that is also differentially expressed in poor outcome groups of patients treated with N-BPs. We functionally validated this patient variation in *ATRAID* as conferring cellular hypersensitivity to N-BPs. Our work adds key insight into the mechanistic action of N-BPs and the processes that might underlie differential responsiveness to N-BPs in people.

**One Sentence Summary:** *ATRAID* is essential for responses to the commonly prescribed osteoporosis drugs nitrogen-containing bisphosphonates.

**Overline:** BONE

## INTRODUCTION

Nitrogen-containing bisphosphonates (N-BPs) are the standard treatment for osteoporosis and several other bone diseases (*1, 2*). Certain N-BPs (pamidronate, zoledronate) are also routinely prescribed to prevent skeletal complications in patients with multiple myeloma and with bone metastases from other malignancies, including breast and prostate cancer (*3*). However, because N-BPs cause rare yet serious side-effects, such as atypical fractures (AFFs) and osteonecrosis of the jaw (ONJ), many patients avoid taking them (*1, 4-6*), causing the number of prescriptions to plummet over 50% in the last decade (*6, 7*). A plan for addressing this crisis, developed by American Society for Bone and Mineral Research (ASBMR) leadership, calls for better pharmacogenomics to identify genetic factors that may underlie response to this class of drugs (*6*).

A goal of personalized medicine is to identify biomarkers that underlie drug responsiveness. For N-BPs, it can be said that there are limited personalization options owing to the limited number of genes implicated in the pharmacologic effects of N-BPs. Exposure of cells to N-BPs leads to inhibition of farnesyl diphosphate synthase (FDPS, also known as FPPS) resulting in reduction in protein prenylation (*8*). On the basis of this observation, it is widely believed that N-BPs act therapeutically by impairing protein prenylation, ultimately leading to deficits in numerous cellular processes including differentiation, recruitment, and adhesion of osteoclasts (the major bone resorptive cell type) to bone and/or osteoclast cell death (*9-11*).

Recently we performed CRISPRi-based, genome-wide screening and identified a poorly characterized gene, *SLC37A3*, that provides molecular details for how N-BPs reach their target, FDPS (*12*). As part of that work we determined that SLC37A3 requires another poorly characterized protein, ATRAID, for its expression (*12*). Here, we independently identified *ATRAID* using a different genome-wide, mutagenesis strategy. We generated *Atraid*-deficient mice and determined that it is required for the regulation of N-BPs on bone. We also performed exome sequencing in patients taking N-BPs and identified and functionally validated rare coding variants in *ATRAID* in patients that suffered side effects, namely atypical femoral fractures and osteonecrosis of the jaw.

## RESULTS

### *ATRAID* is required for molecular responses to nitrogen-containing bisphosphonates

To provide insight into the mechanism(s) of N-BPs action, we performed a genetic screen to identify human genes required for the anti-proliferative effects of N-BPs (Fig. 1A). We used a largely haploid human cell line of myeloid origin (KBM7, also known as HAP1) to generate a library of retroviral gene trap mutants (*13*) and then selected for clones that are resistant to cytotoxic concentrations of alendronate (ALN). The advantages of this cell line for genetic screening include: i) each gene is present as a single copy, enabling gene inactivation (except those genes on chromosome 8); ii) KBM7 cells are human and of the hematopoietic lineage, increasing the likelihood that any genes we identify could be relevant to the natural context for N-BPs, the bones of human patients (*14*); and iii) it is a different cell line and a different mutagenic approach than used previously with CRISPRi in K562 cells (*12*), which allows us to independently assess those results. Using this haploid approach, we identified, *ATRAID*, also known as *APR-3/C2orf28* (*15*), as the gene most significantly enriched for insertions in alendronate-resistant cells compared to untreated cells (FDR corrected *P*-value = 7.02e-45) (Fig. 1B; fig. S1, A and B, and table S1). Providing confidence in our screen, we also identified *SLC37A3*, as well as *SNTG1, PLCL1*, and *EPHB1*, which have been previously connected to N-BP action on bone cells and/or human bone diseases (table S1) (*16-18*).

**Fig. 1.**
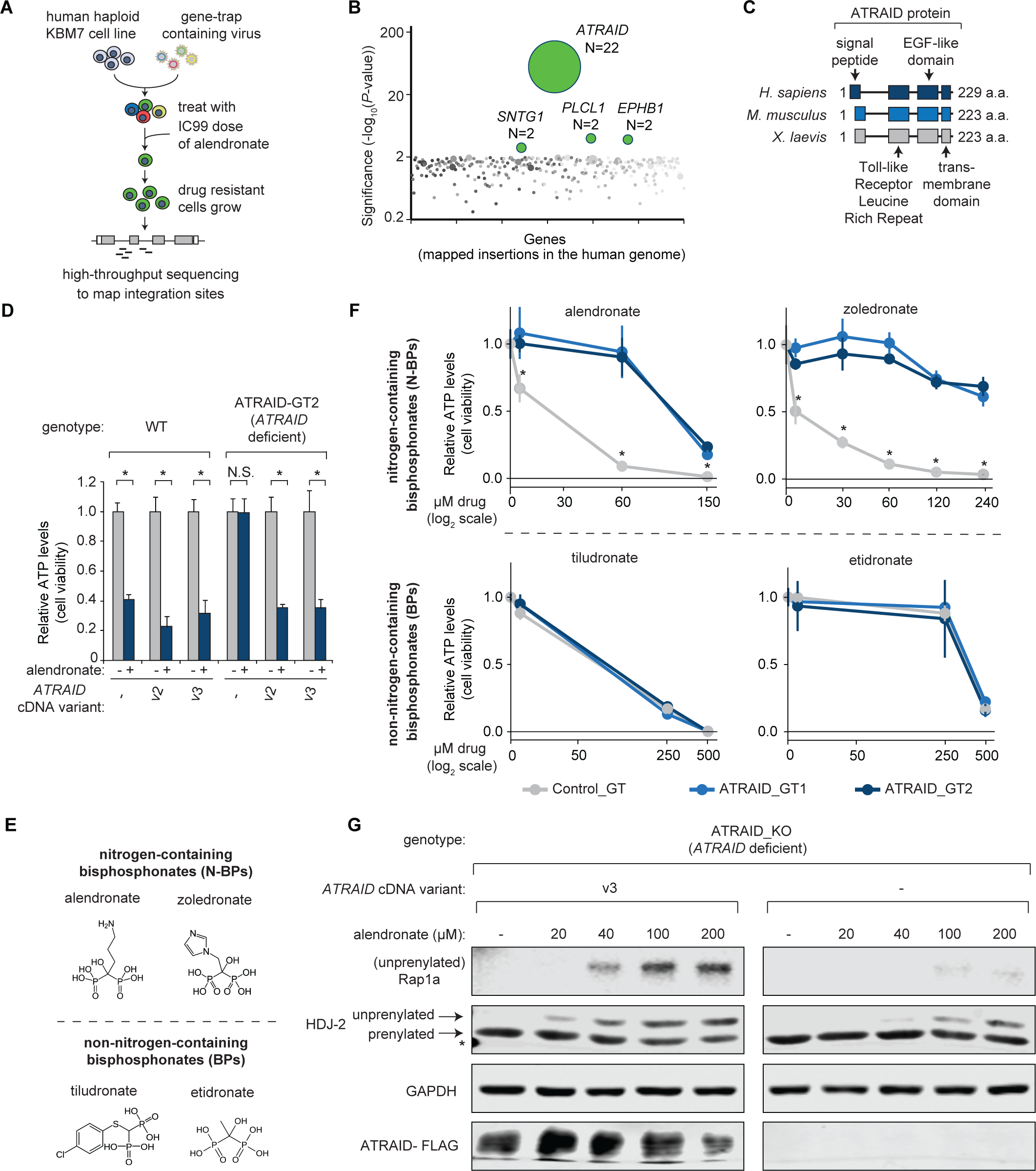
*ATRAID* is required for molecular responses to nitrogen-containing bisphosphonates. **(A)** Schematic of haploid mutagenesis screening pipeline. Sequencing-based identification of gene-trap insertion sites in alendronate-resistant human haploid KBM7 cells. Genomic DNA for sequencing was obtained from mutagenized KBM7 cells grown for four weeks post-treatment with alendronate (165 µM). **(B)** Sequencing-based identification of gene-trap insertion sites in alendronate-resistant cells. N=number of unique insertions within the stated gene locus. False discovery rate corrected (FDR) *P*-values for *ATRAID*=7.02×10^−45^, *PLCL1*=1.02×10^−04^, *EPHB1*=2.05×10^−04^, *SNTG1*= 1.84×10^−03^. *P*-values represent enrichment in alendronate-treated versus vehicle treated cells. **(C)** Schematic representation of structural features of human ATRAID protein and its mouse and frog orthologues. **(D)** Cell viability in wild-type control and *ATRAID*-deficient cells exogenously expressing or not expressing *ATRAID* cDNA. Cells were treated with alendronate (60 µM) and analyzed for cell viability. Cell viability was determined by measuring cellular ATP and is expressed as a ratio of that compared with untreated cells. Error bars indicate the standard deviation for n=4 (biological replicates). N.S., not significant; \**P* < 0.05, student’s *t-*test. v2, variant 2 (NM_080592.3); v3, variant 3 (NM_001170795.1) of the *ATRAID* gene, respectively. **(E)** Chemical structures for nitrogen-containing bisphosphonates (N-BPs) or non-nitrogen-containing bisphosphonates (BP). KBM7 cell viability in *ATRAID*-deficient (ATRAID_GT1 and ATRAID_GT2) and control (wild-type) KBM7 cells upon treatment with nitrogen-containing bisphosphonates (N-BPs) or non-nitrogen-containing bisphosphonates (BP). All cells were treated with the indicated concentration of the indicated N-BP (alendronate, zoledronate), BP (etidronate, tiludronate) for 72 hours. Cell viability was determined by measuring cellular ATP and is expressed as a ratio of that compared with untreated cells. All measurements were performed in quadruplicate (biological replicates). \**P* < 0.05, student’s *t*-test. **(F)** Immunoblots of cell lysates from *ATRAID*-deficient and *ATRAID* v3-reconstituted HEK-293T cells treated with the indicated dose of alendronate for 24 hours. Equal amounts of protein were loaded in each lane. This experiment was repeated three times (biological replicates) and was consistent all three times. *non-specific band.

*ATRAID* was named because it is a gene whose mRNA expression is strongly induced by the ligand all-trans retinoic acid (*15*). *ATRAID* is conserved in chordates and contains a signal peptide, Toll-like-receptor leucine rich repeat, EGF-like domain, and a transmembrane domain (Fig. 1C and fig. S1A) (*19, 20*). The alendronate resistance phenotype of *ATRAID*-deficient cells (ATRAID_GT1 (gene-trap1) and ATRAID_GT2 (gene-trap2) was reversed by the re-introduction of wild-type *ATRAID* splice variant 2 (v2) or splice variant 3 (v3) cDNA, which differ in their N-termini (Fig. 1D; fig. S1, A and B). To better understand the degree of alendronate resistance in *ATRAID*-deficient cells, we varied both cell number and drug concentration in the viability assay. *ATRAID*-deficient cells were resistant to alendronate over two to three orders of magnitude of drug concentration or cell number (fig. S1C). The growth of untreated wild-type and *ATRAID*-deficient cells didn’t differ (fig. S1D). Overexpression of full-length *ATRAID* (v2) sensitized cells to alendronate (fig. S1E). Lastly, ATRAID membrane targeting is required for the anti-proliferative effects of alendronate, as *ATRAID-*deficient cells complemented with full-length *ATRAID* (v2) were sensitive to the cytotoxic effects of alendronate, whereas those expressing the membrane truncated form remained resistant (fig. S1F). Taken together, these data establish *ATRAID* as a genetic factor required for the growth inhibitory effects of alendronate.

N-BPs are part of a larger class of compounds known as bisphosphonates (BPs) that contain two phosphate moieties each joined to a carbon atom by a carbon-phosphorus bond (*21*) (Fig. 1E). To determine whether the effects of *ATRAID* deficiency on alendronate resistance were specific to nitrogen-containing bisphosphonates, we tested the effect of several nitrogen-containing and non-nitrogen-containing bisphosphonates on wild-type and mutant *ATRAID* cells. *ATRAID*-deficient cells were resistant to the nitrogen-containing bisphosphonates, alendronate and zoledronate (ZOL), but were as sensitive to the non-nitrogen-containing bisphosphonates, etidronate and tiludronate, as control cells (Fig. 1E).

To determine whether *ATRAID* is required for the reduction in protein prenylation observed upon N-BP treatment, we monitored the prenylation of several proteins, including the heat shock protein DnaJ (Hsp40) homolog HDJ-2, and the Ras family GTPase Rap1a (*22*). Alendronate strongly inhibited prenylation of HDJ-2 and Rap1a in wild-type cells in a dose dependent manner and had much less of an effect on prenylation of these proteins in *ATRAID-*deficient cells (Fig. 1F). Furthermore, the inhibitory effect of alendronate on prenylation was rescued when *ATRAID* cDNA variants (v2 and v3) were introduced (fig. S1F). We observed inhibition of prenylation resistance at N-BP doses where we did not see PARP-1 cleavage in *ATRAID* deficient cells, suggesting that *ATRAID* can mediate the effect on prenylation independent of apoptosis (fig. S1G). Thus, these findings suggest *ATRAID* functions as a positive regulator upstream of FDPS.

### ATRAID is required for organismal responses to nitrogen-containing bisphosphonates

To determine whether ATRAID modulates organismal responses to N-BPs we inactivated *Atraid* globally in mice (*23*). We confirmed deletion of *Atraid* exons 3-5 and determined that *Atraid* homozygous deleted *Atraid*^KO^ mice (labeled KO, -/-) are viable but their body weight is mildly reduced compared with litter-matched derived, wild-type controls (labeled as WT, +/+) (fig. S2, A to C). We confirmed that tail fibroblasts derived from *Atraid*^KO^ animals are resistant to the cytotoxic effects of alendronate (fig. S2, D and E). Before studying the effects of the N-BPs in the context of *Atraid* loss, we first characterized the basal role of *Atraid* in bones. We determined that *Atraid* mRNA expression was undetectable in the bones of *Atraid*^KO^ animals and that *Atraid*^KO^ mice had slightly smaller bones compared with litter-matched derived, wild-type control mice (fig. S2, F and G). To examine the effects of *Atraid* on bone structure, we performed micro-computed tomography (µCT) analysis (*24*). *Atraid* deficiency did not decrease either trabecular or cortical structural parameters (fig. S2, H and I; data file S1). We measured bone strength using three-point bending tests (*25*). Some measures, such as stiffness (Newtons/millimeter, N/mm) and post-yield displacement (a measure of bone fragility, in millimeters, mm) were decreased by *Atraid* deficiency, whereas others, such as yield load (the point where bone bending goes from elastic vs. plastic, in Newtons, N), were not significantly altered (fig. S2, J to L; data file S1, *P* > 0.05, student’s *t-*test).

Osteoclasts release degradation products from C-terminal telopeptides of type I collagen (CTX-I) from bone into blood (*26*), and CTX-I in serum was not significantly different in wild-type mice compared with *Atraid*^KO^ mice (fig. S2M; data file S1, *P* > 0.05, student’s *t-*test). Histomorphometric measures of osteoclast function including osteoclast surface per bone surface (Oc.S/BS) (*27*), as judged by Tartrate Resistant Acid Phosphatase (TRAP) staining (*28*), were also not statistically different (fig. S2N; data file S2). Consistent with a basal defect in osteoblast function (*29*), *Atraid*^KO^ mice have reduced serum circulating Gla-Osteocalcin [Gla-OC; the activated form of osteocalcin, incorporated in bone matrix (*30*)] and modestly reduced bone formation rate (BFR) as measured by double-labeling (*27*) (fig. S2, O and P; data file S1).

To test the effect of alendronate in a model that mimics menopausal bone loss, the most common indication for the N-BPs, we performed ovariectomies (OVX) on adult female mice (Fig. 2A) (*31*). When ovaries are removed from females, the changes in estrogen cause a reduction in bone density triggered by disruption of the balance of osteoblast and osteoclast functions. This loss of bone density can be alleviated by treatment with N-BPs (*32*). The magnitude of trabecular bone loss in WT and *Atraid*^KO^ sham mice four weeks after OVX is exemplified in the µCT 3D reconstruction of the femoral proximal metaphysis (Fig. 2B). Consistent with alendronate preventing bone loss (*32*), femoral cortical and trabecular structural parameters, including cortical thickness and area, bone volume/trabecular volume (%), and trabecular thickness, were increased by alendronate treatment of WT OVX mice (Fig. 2, C to F and data file S1; see WT OVX +/- alendronate). In contrast, alendronate had blunted effects in *Atraid*^KO^ OVX mice (Fig. 2, C to F and data file S1; see *ATRAID*^KO^ OVX +/- alendronate). The same patterns of alendronate resistance in *Atraid*^KO^ mice were observed for bone strength (Fig. 2, G and H). That is, alendronate increased bone strength as judged by stiffness and yield load in wild-type ovariectomized mice, but its effects were blunted in *Atraid*^KO^ matched cohorts (Fig. 2, G and H; data file S1). Taken together, these results suggest that *Atraid* is required for the beneficial effects of N-BPs in ovariectomized female mice.

**Fig. 2.**
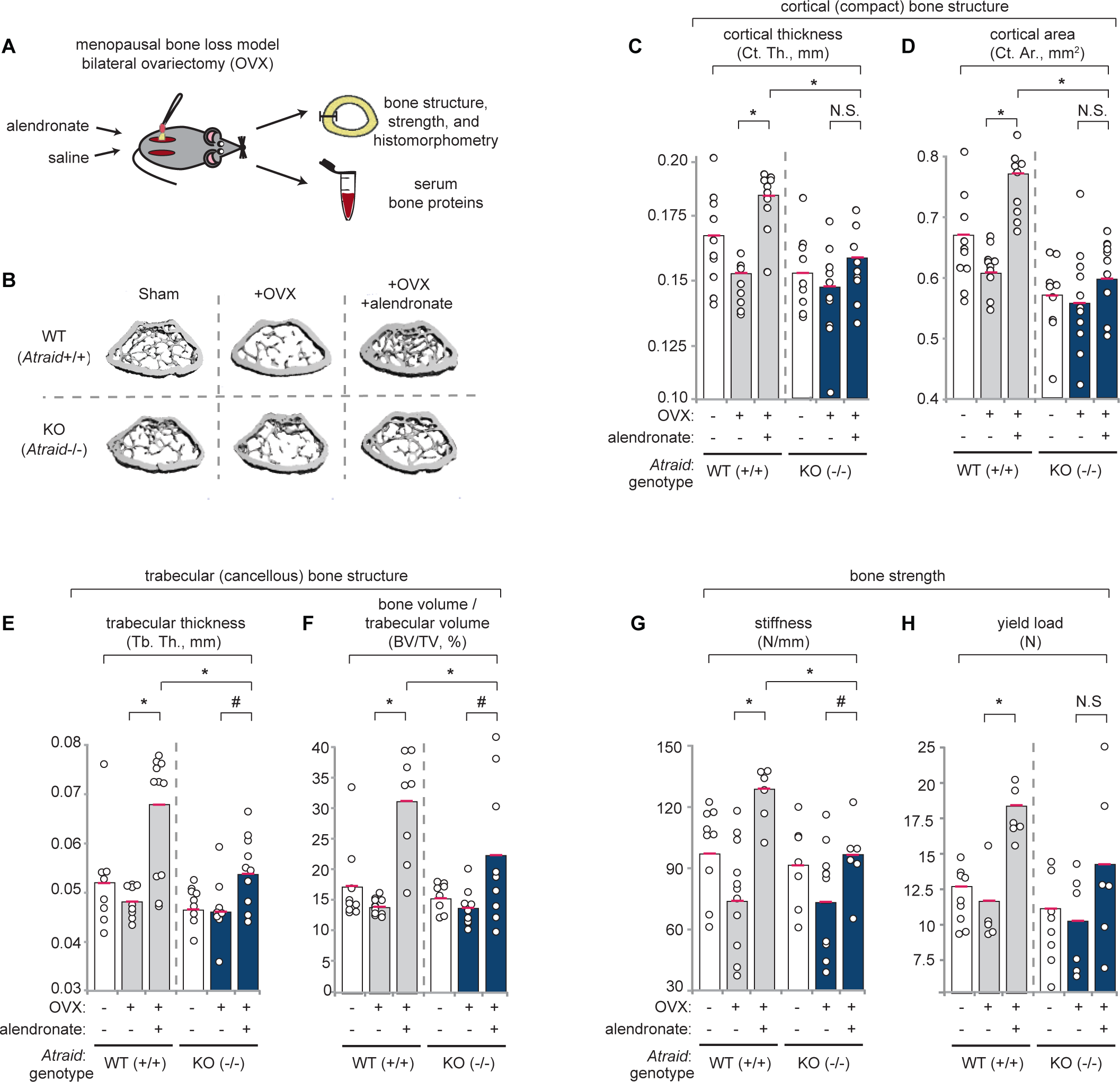
*Atraid* is required for organismal responses to nitrogen-containing bisphosphonates. **(A)** Schematic of mouse menopausal bone loss model – bilateral ovariectomy (OVX). Saline or 100 μg/kg/week alendronate was administered concurrent with OVX or a sham procedure. After four weeks, mice were euthanized and bones and serum were extracted and analyzed. **(B)** Representative µCT reconstructions of femoral trabecular bone from 4-month-old litter-matched derived, wild-type, *Atraid* WT (+/+), and KO (-/-), female mice that were either ovariectomized (OVX) or sham operated (Sham), treated with either vehicle (saline) (+OVX), or alendronate (+OVX+ALN) for four weeks. **(C-F)** Ovariectomized WT and *Atraid* KO mice and their bone microstructural responses to alendronate. Femur cortical (C, D) and trabeculae (E, F) regions were analyzed by µCT. Each circle represents an individual animal. Circles offset to the right represent unique animals with similar values to those of another animal (offset for visual clarity). N=6-11 mice (3.5 month old) per group. \**P* < 0.01, ^#^ indicates 0.01<*P*<0.05, student’s *t*-test and red line indicates mean. **(G-H)** Ovariectomized WT and *Atraid* KO mice and their bone strength responses to alendronate. Stiffness (G) and yield load (H) were analyzed by three-point bending test. Each circle represents an individual animal. Circles offset to the right represent unique animals with similar values to those of another animal (offset for visual clarity). N=6-11 mice per group. \**P* < 0.01, ^#^0.01 < *P* < 0.05, N.S., indicates not significant, student’s *t*-test, and red line indicates mean.

To test an additional osteoporosis model, we examined senile (old age) osteoporosis using 18 month-old male WT and *Atraid* deficient mice (*33*). After treating these mice weekly with alendronate or saline for two months, we found similar results to those in our OVX study. That is, measures of bone density were increased by alendronate, but less so in the *Atraid* deficient mice (fig. S2, Q and R, data file S1). This further suggests *Atraid* is key for responses to N-BPs in vivo.

### *ATRAID* is required cell-autonomously for N-BP inhibition of osteoclast function

Because N-BPs potentially affect osteoclasts and osteoblasts, we investigated whether *Atraid* deficiency would regulate the effects of alendronate in each cell type in our post-menopausal (OVX) and old-age (senile) osteoporosis models. Regarding osteoclasts, in wild-type mice both serum and bone histological markers of osteoclast function, CTX-I, and osteoclast surface per bone surface (Oc.S/BS) and osteoclast number per bone surface (N.Oc/BS), respectively, were impaired by alendronate treatment (Fig. 3, A and B; fig. S3, A and B; data file S2) in both osteoporosis models. In contrast, in *Atraid*^KO^ mice, alendronate was less effective on osteoclasts in both osteoporosis models (Fig. 3, A and B; fig. S3, A and B; data file S2). That osteoclast number was reduced by N-BPs in wild-type mice is consistent with our cell viability measurements in non-osteoclasts and with previous literature (*32*).

**Fig. 3.**
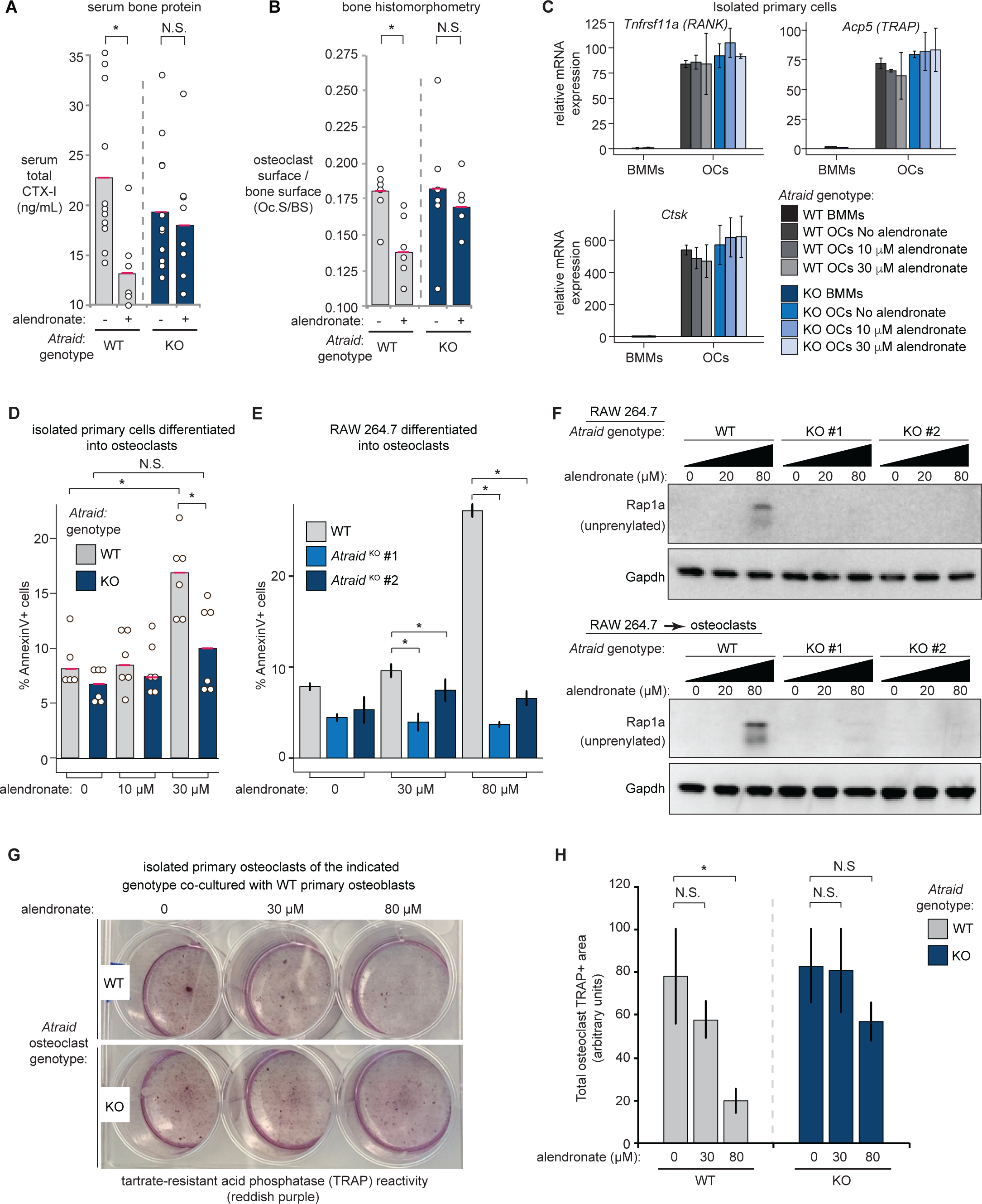
*Atraid* is required cell-autonomously for N-BP inhibition of osteoclast prenylation. **(A)** CTX-I, a serum marker of osteoclast activity, was measured in WT and *Atraid*^*KO*^ ovariectomized mice with or without alendronate treatment by ELISA. Each circle represents an individual animal. Circles offset to the right represent unique animals with similar values to those of another animal (offset for visual clarity). N=8-13 mice per group. \**P* < 0.05, student’s *t*-test. **(B)** Osteoclast histomorphometric responses in WT and *Atraid*^*KO*^ ovariectomized mice with or without alendronate treatment. Osteoclast surface to bone surface ratio (Oc.S/BS) was determined by Tartrate Acid Phosphatase (TRAP)-assay reactivity. Each circle represents an individual animal. Circles offset to the right represent unique animals with similar values to those of another animal (offset for visual clarity). N=5-7 mice per group. \**P* < 0.05, n.s. indicates not significant, student’s *t*-test, and red line indicates mean. **(C)** Quantitative PCR to examine mRNA expression of markers of osteoclast differentiation, *Ctsk, Tnfrsf11a (RANK), Acp5 (TRAP)*, in wild-type (WT) and *Atraid*^KO^ M-CSF-expanded bone marrow macrophages (BMMs) differentiated with *RANKL* to osteoclasts. Expression is normalized to wild-type, undifferentiated BMM cells, using *Actb* and *Rplp0* as control genes. Error bars represent the standard deviation of technical triplicate reactions. **(D)** Percent of Annexin-V positive cells after a 48 hour alendronate treatment of WT and *Atraid*^KO^ BMMs differentiated into osteoclasts. Annexin V staining was assessed using flow cytometry. Each circle represents osteoclasts derived from an individual animal (split for treatment with 0, 10 μM, 30 μM alendronate). Red line indicates mean. \**P* < 0.05, N.S. indicates not significant, student’s *t-*test. **(E)** Percent of Annexin-V positive cells after a 48 hour alendronate treatment (0, 30 μM, 80 μM) in wild-type and *Atraid*^KO^ differentiated RAW 264.7 osteoclasts. Annexin V staining was assessed using flow cytometry. Error bars represent the standard deviation of n=3 experiments (biological replicates), \**P* < 0.05, student’s *t-*test. **(F)** Immunoblots of cell lysates of RAW wild-type (WT) and *Atraid*^KO^ (KO) cells, and RAW 264.7-derived osteoclasts treated with alendronate for 48 hours. Top panel: immunoblot specific to the unprenylated version of Rap1a. Bottom panel: Gapdh, serving as a loading control. Alendronate concentrations were 0, 20 μM, 80 μM. **(G)** Representative image of a six-well dish co-culture of equal numbers of mouse primary osteoblasts and osteoclasts of the indicated genotypes with or without the indicated doses of alendronate (ALN) for four days. The experiment was performed three independent times with a similar result. Red staining reflectsTRAP-assay reactivity. **(H)** Image analysis of the samples in (G). Error bars represent the standard deviation of n=3 independent images (technical replicates). \**P* < 0.01, N.S. indicates not significant, student’s *t-*test.

To provide insight into the effects of N-BPs on osteoblasts in our osteoporosis models, we measured BFR and mineral apposition rate (MAR) (*27*). Unlike BFR in which the rate is normalized by the amount of labeled bone surface, MAR is the rate of bone formation irrespective of how much of the bone is active (*27*). Alendronate did not affect trabecular MAR or BFR in either wild-type or *Atraid*^KO^ mice in either osteoporosis model (fig. S3, C and D; data file S2).

To determine whether *Atraid* is required in a cell autonomous manner for the N-BP-dependent effects on osteoclasts, we isolated M-CSF-expanded bone marrow macrophages (BMMs) from both WT and *Atraid*^KO^ mice, and differentiated these cells into osteoclasts following a standard protocol (*34*). *Atraid*^KO^ BMMs differentiated into osteoclasts as well as wild-type cells irrespective of treatment with alendronate, yet BMM-derived *Atraid*^KO^ osteoclasts were resistant to alendronate-induced apoptosis (Fig. 3, C and D). This suggests that *Atraid* is required cell autonomously in osteoclasts for the effects of N-BPs on cell number.

As an independent confirmation of our primary cell experiments, we generated *Atraid* knockout RAW 264.7 cells and differentiated them to osteoclasts (fig. S3E). RAW 264.7 cells are a robust, well-characterized murine macrophage cell line that can be differentiated to osteoclast-like cells using RANKL (*35*). We treated both RAW 264.7 cells and the RAW 264.7 cells differentiated into osteoclasts with alendronate, and found *Atraid* deficiency, as expected, conferred resistance to doses that induced apoptosis (Fig. 3E). Alendronate did not affect known markers of osteoclast differentiation in wild-type cells (Fig. 3C). Therefore, to pursue the mechanism of N-BPs on osteoclast function, we focused on prenylation. In alendronate-treated RAW 264.7 cells and osteoclasts differentiated from RAW 264.7 cells, we found that *Atraid*^KO^ cells were resistant to alendronate-induced inhibition of prenylation (Fig. 3F).

We assessed whether wild-type osteoblasts might sensitize *Atraid*-deficient osteoclasts to N-BPs. We cultured primary wild-type osteoblasts with either WT or *Atraid*-deficient primary osteoclasts and treated these co-cultures with alendronate or vehicle. As in the case of WT RAW 264.7 cells grown independently (Fig. 3, E and F), WT osteoclasts were more inhibited by alendronate than *Atraid* deficient osteoclasts despite the presence of WT osteoblasts in both cases (Fig. 3, G and H). In total, these findings support that *Atraid* is required for the cell-autonomous effects of N-BPs on osteoclasts.

### Genetic factors involved in responses to nitrogen-containing bisphosphonates in patients

We sought an unbiased approach to determine what genes might be relevant in patients treated with N-BPs. We performed whole exome sequencing (WES) on two sets of patients taking N-BPs who experienced side effects: patients with osteoporosis who experienced atypical femoral fractures (AFF) (n = 27 patients), as well as patients with multiple myeloma or breast cancer patients who experienced osteonecrosis of the jaw (ONJ) (n = 8 patients) and 11 control patients taking N-BPs that didn’t experience AFF or ONJ (Fig. 4A and data file S3 for patient information). We also analyzed two published gene expression datasets involving patients who had taken N-BPs: patients with multiple myeloma who did or did not suffer ONJ when taking N-BPs (*36*), and patients with breast cancer with bone marrow disseminated tumor cells (DTC) which reoccurred or the patient died less than 1000 days vs. greater than 2500 days following initiation of zoledronate treatment (*37*) (Fig. 4A and data file S3). We then compared the patient data to three cell-based, genome-wide CRISPRi and CRISPRa screens we previously performed: alendronate and zoledronate CRISPRi and alendronate CRISPRa (*12, 38*) (data file S3). To identify genes involved in N-BP response across experimental paradigms we generated a Venn diagram to visualize the overlap of “hits”. In comparing the WES hits –genes that had the same rare coding variants (minor allele frequency < 0.05) in both patients with AFF and ONJ but not in controls – with our hits from our alendronate and zoledronate CRISPRi/a screens, we identified 64 genes in common including *ATRAID, FDPS*, and *SLC37A3* (Fig. 4A). When comparing the WES hits with the gene expression hits, we identified 49 genes, whereas the CRISPRi/a and gene expression studies had 76 genes in common (Fig. 4A). Three genes, *ATRAID, ATR*, and *ZBTB4* were statistically significant in all three data types (Fig. 4A) (data file S3, FDR corrected *P* < 0.05). Focusing on *ATRAID* specifically, we observed a ∼50% decrease in *ATRAID* expression in the patients with DTC and ONJ that had their gene expression measured. By exome sequencing the patients with AFF and ONJ, in *ATRAID* we detected two rare variants – hereafter referred to as the ‘D5G/G32R variant’ – that were present together in 3 out of 35 patients with AFF and ONJ (2 out of 27 AFF; 1 out of 8 ONJ) (Fig. 4B).

**Fig. 4.**
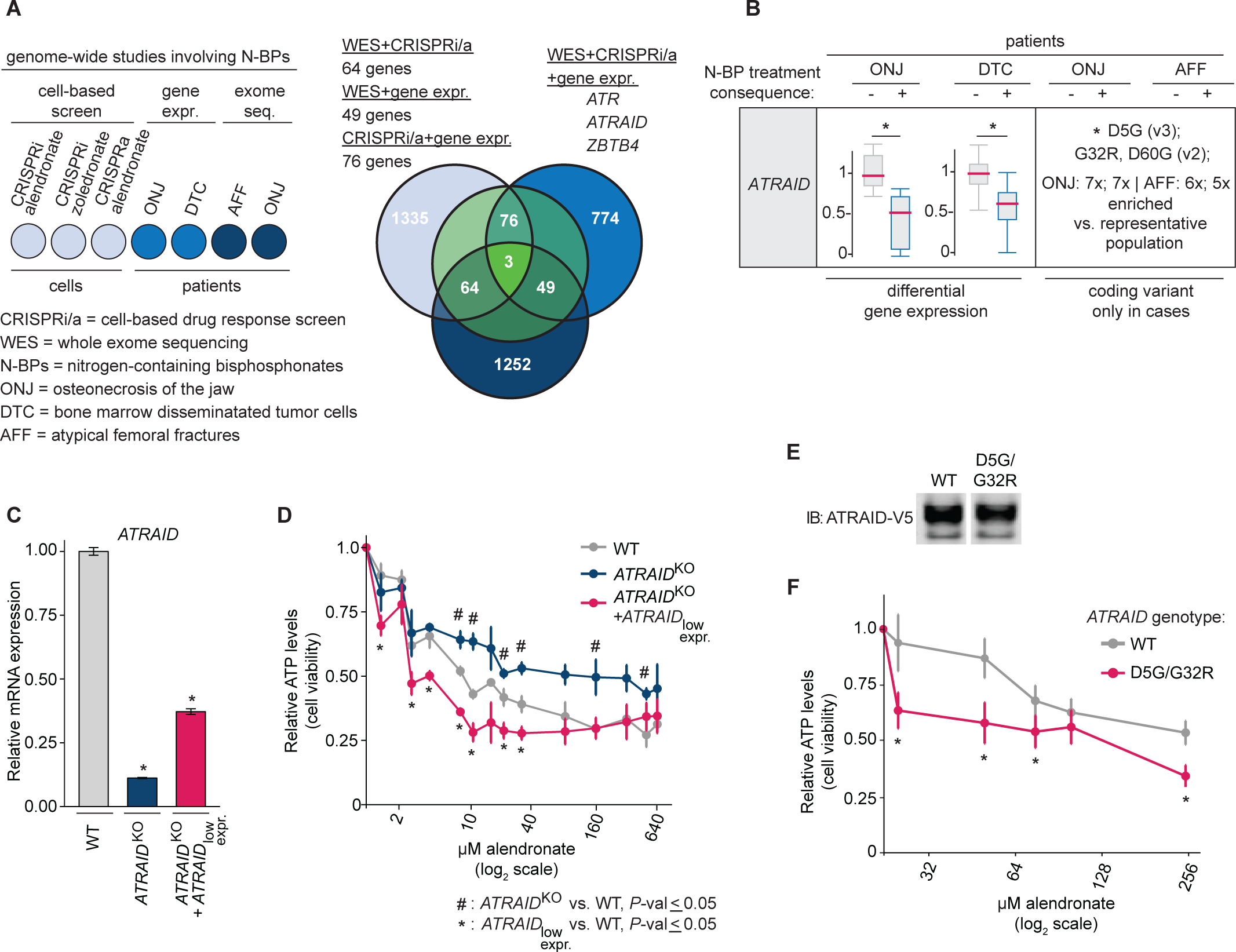
*ATRAID* as a potential genetic factor for altered responses to nitrogen-containing bisphosphonates in patients. **(A)** Genome-wide studies of N-BP responsiveness in patients vs. cells. The outcomes considered from human studies involving N-BPs are: osteonecrosis of the jaw (ONJ), breast cancer bone marrow micrometastases [disseminated tumor cells (DTC)], and atypical femoral fractures (AFF). *ATRAID, ATR, ZBTB4* are statistically significant hits (FDR corrected *P* < 0.05) in N-BP cell-based CRISPRi/a screening, differentially expressed in both gene expression datasets (ONJ and DTC), and possess rare multiple non-synonymous coding variants in AFF and ONJ cases but not controls. These three genes are visualized as the Venn diagram of overlap of lists of genes that met the following criteria: significant alendronate CRISPRi, zoledronate CRISPRi, or alendronate CRISPRa hits with absolute value of rho growth phenotype values ≥ 0.30 and *P* ≤ 0.05 (1335 out of 15828 genes) (*12, 38*); differentially expressed N-BP in ONJ + DTC (774 out of 18415 for ONJ (*36*) and 20492 for DTC (*37*); multiple coding variant(s) in AFF and ONJ cases and not controls (1252 out of 11659 genes) (data as part of this study). **(B)** Patient genetic data for *ATRAID*. Raw expression values were normalized to 1 to fit on a comparable Y axis. \**P* < 0.05, moderated *t*-test. N.S. indicates not significant. “X”-enriched refers to the fold-enrichment of the allele compared with a population with a similar genetic background as the cases. For example, for *ATRAID*, the D5G variant is present in 2 out of 27 AFF patients. Though this allele wasn’t detected in the 11 control samples, it is present in a population of European Americans (EA) and Asian Americans (AA) that is representative of the study population at a prevalence of 0.0131. Therefore, the D5G allele is (2/27) / 0.0131 = 5.66X enriched in cases compared to the EA/AA population. “v3” and “v2” refer to isoforms of the *ATRAID* gene. A simple binomial test was used to calculate the significance of each variant. * *P* < 0.05. **(C)** Quantitative PCR to examine *ATRAID* mRNA expression in wild-type, ATRAID-deficient, and low ATRAID expressing cells. Error bars represent the standard deviation of technical triplicate reactions. Expression was normalized to WT cells using *RPLP0* and *TBP* as controls. * indicates *P*<0.05, student’s *t-*test. **(D)** Cell viability in wild-type, *ATRAID*-deficient, and low *ATRAID* expressing cells. Cells were treated with the indicated doses of alendronate and analyzed for cell viability. Cell viability was determined by measuring cellular ATP and is expressed as a ratio of that compared with untreated cells. Error bars indicate the standard deviation for n=4 (biological replicates). N.S., not significant; * and ^#^ indicate *P*<0.05 for the indicated cell lines, student’s *t-*test. **(E)** Immunoblot (IB) of wild-type and D5G/G32R variant *ATRAID*-V5 tagged proteins. Mutant or wild-type *ATRAID* v2 and v3 were stably introduced into *ATRAID*-deficient HEK-293T cells. **(F)** Cell viability in wild-type vs. D5G/G32R variant cells. Cells were treated with the indicated doses of alendronate and analyzed for cell viability. Cell viability was determined by measuring cellular ATP and is expressed as a ratio of that compared with untreated cells. Error bars indicate the standard deviation for n=4 (biological replicates). N.S., not significant; \**P* < 0.05, student’s *t*-test.

We sought to determine the functional relevance of decreased *ATRAID* mRNA expression and the *ATRAID* D5G/G32R variant, both of which are associated with bad outcomes of N-BP treatment (Fig. 4, A and B). To test the former, we expressed *ATRAID* mRNA at sub-endogenous quantities in *ATRAID* deficient cells such that the expression was similar to the reduced expression seen in patients that experienced ONJ or DTC (∼50% compared to wild-type controls) (Fig. 4C). We refer to these *ATRAID* partially restored cells as ‘*ATRAID*_low expr._’. *ATRAID*-deficient cells conferred resistance to alendronate as expected, whereas *ATRAID*_low expr._ cells were hypersensitized to alendronate (Fig. 4D). Similarly, the *ATRAID* D5G/G32R variant, which we identified in both the patients with AFF and ONJ, conferred hypersensitivity to alendronate compared with wild-type *ATRAID* (Fig. 4, E and F). Taken together, this suggests that bad patient outcomes might reflect cellular hyper-response to N-BPs. In total, these findings support the importance of ATRAID in bisphosphonate responsiveness in humans.

## DISCUSSION

This work focused on the physiological impact of *ATRAID* as a positive regulator genetically upstream of *FDPS*. Here we use prenylation as an output of FDPS function. Recently, we linked *FDPS* to DNA synthesis and damage (*39*). This was intriguing in light of earlier studies in the context of ONJ where N-BPs regulated p63 – a well-known mediator of DNA damage (*40*) – in a mevalonate pathway-dependent manner (*41*). Considering that each of the three top genes from the patient analysis, *ATRAID, ATR*, and *ZBTB4*, are involved in p53 responses – a better known p63-related mediator of DNA damage (*42-45*) – it will be interesting to determine whether these genes mediate their effects on p53/p63 signaling via *FDPS*.

The molecular effects of N-BPs on FDPS require the transporter, SLC37A3 (*12*). Interestingly, the SLC37A family member (*46*), *SLC37A2*, is mutated in dogs and gives rise to a bone overgrowth phenotype that resembles the human disease Caffey syndrome (*47, 48*). This phenotype is particularly interesting because it suggests that natural ligands or drugs that inhibit the SLC37A family might phenocopy N-BP treatment in increasing bone density.

ATRAID binds NELL-1, a secreted protein that promotes bone mineralization in mice and potentiates osteoblast differentiation in an *ATRAID*-dependent manner (*49, 50*). It is also notable that NELL-1 is in pre-clinical testing for the treatment of osteoporosis (*51*). In future studies, it will be interesting to determine whether NELL-1 affects the responses to N-BPs we observe upon manipulating *ATRAID*. NELL-1 has a related family member, NELL-2. This family member has been the subject of high profile studies in the field of axon guidance (*52*). It is unknown whether ATRAID signals to NELL-2 and if so what role it may play in the brain.

There are several limitations of our study. 1) Because we used a global knockout strategy with *Atraid*, we can’t definitively conclude it is required in vivo in osteoclasts – the target cell type for the N-BPs; 2) We identified *ATRAID* in screening in leukemia cells, not in osteoclasts.

Therefore, there it is possible a screen in a cell type more relevant to the N-BPs would yield additional genes important to bone; 3) There are relatively modest numbers of DNA samples in existence from ONJ and AFF patients. More samples need to be collected and analyzed to further test our findings as identifying those patients who might experience these consequences when taking N-BPs is of paramount importance.

## MATERIALS AND METHODS

### Study design

The objective of this study was to identify and subsequently characterize factors involved in the on- and off-target effects of the osteoporosis drugs, nitrogen-containing bisphosphonates (N-BPs). To address this, we performed a genome-wide haploid cell screen and identified the gene *ATRAID*. To assess the cellular role of ATRAID in the response to N-BPs, we treated a variety of cell lines including human 293T and KBM7 cells, murine macrophage RAW 264.7 cells differentiated into osteoclasts and primary cells derived from mice, with the N-BPs (including alendronate and zoledronate) or other drugs and assessed cell viability/growth/fitness by measuring cellular ATP, as well as protein prenylation by immunoblot. These analyses established that ATRAID is required for the cellular responses to N-BPs. We investigated the in vivo role of ATRAID by utilizing two mouse models of osteoporosis (ovariectomies on 3.5-month old female mice as a model of post-menopausal osteoporosis (OVX), and 18-month old male mice as a model for senile osteoporosis) treated with alendronate. The effects on wildtype and *Atraid*^KO^ mice were assessed by profiling bone structure using micro-computed tomography (µCT) (OVX, senile: n = 6-11, n = 5-8), strength using a three-point bending assay (OVX: n = 6-11), histomorphometry using TRAP staining and double-labeling (OVX, senile: n = 5-7, n = 4-7), and serum bone proteins using ELISA (OVX, n = 8-13). These analyses established that ATRAID is required for the organismal responses to N-BPs. To translate our findings to humans, we integrated clinical and unbiased genome-scale molecular data in patients treated with N-BPs, including patients that experienced atypical femoral fractures or osteonecrosis of the jaw while being treated with N-BPs. These analyses established that ATRAID is potentially important for N-BP responses in humans. For our animal studies, mice were randomized to treatment groups, and subsequent analyses were blinded to the extent possible. All experiments involving mice were performed with protocols approved by the Harvard and Washington University Animal Studies Committees. The details of study design, sample sizes, experimental replicates, and statistics are provided in the corresponding figures, figure legends, data files, and Material and Methods.

### Statistical analysis

Unless otherwise specified, group means were compared by one-tailed student’s *t* test for unpaired samples. Data on repeated measures were analyzed by ANOVA, followed by a post-hoc multiple Holm–Sidak method t-test. All data are expressed as the mean ± s.d with numbers of samples indicated in figure legends. *P* values are indicated in each figure legend, and values less than 0.05 were considered significant (alpha) with ≥ 80% power, unless indicated otherwise. We estimated the cohort sizes we would need for this study based on our prior study which involved a similar experimental paradigm in using bisphosphonates and the ovariectomy (OVX) osteoporosis model in BL/6 mice (*32*). All code used to generate statistics and correlations for this project can be found at https://github.com/tim-peterson/ATRAID (DOI: 10.5281/zenodo.3739576).

## Supporting information

ATRAID data file S1

ATRAID data file S2

ATRAID Supp. Table 1

ATRAID data file S3

## ACKNOWLEDGMENTS

We thank members of the Peterson and E. O’Shea laboratories for helpful discussions, especially C. Chow and K. Li (Peterson), and A. R. Subramaniam, C. Chidley, and A. Puszynska for illustrations (O’Shea). We thank M. Bouxsein and D. Brooks (Beth Israel Deaconess Medical Center), and D. Lieb, Y. Kim, and M. Silva (Washington University School of Medicine) for bone structure and function analysis. We thank K. Nagano and R. Baron (Harvard School of Dental Medicine), Y. Iwamoto (Massachusetts General Hospital) and D. Novack, G. London, K. Hyrc, and C. Idleburg for histology, histomorphometry, and imaging (Washington University School of Medicine). We thank L. Gilbert (University of California San Francisco) for assistance on the CRISPRi/a screening protocol.

## Author Contributions

T.R.P. and L.E.S. designed the study. D.T.B. performed the co-culture experiments and senile osteoporosis mouse studies. L.E.S. and J.L. performed the *ATRAID* variant studies. J.P., A.R., and Z.Y. performed viability assays. S.K. and L.E.S. performed immunoblots. Y.B., B.H.L, D.T.B. and N.S. conducted analysis of both basal and OVX mouse studies. C.L., F.W., and T.R.P. performed statistical analysis for the patient gene expression and mouse studies. T.R.P. performed the statistical analysis for the CRISPRi/a screens. S.M., J.C.B., M.M., M.S., M.H., S.D., V.N.B., R.C., M.J.G., C.M.M., W.M.R., C.A.G., and K.D. performed the exome sequencing on the AFF patients. G.H. and C.C.G. performed statistical analysis for the exome sequencing of the AFF patients. D.A.H, C.L.C., K.M.S., N.R., T.B.D., and T.R.P. performed the exome sequencing on the ONJ patients. G.H. performed the statistical analysis for the exome sequencing of the ONJ patients. M.V., K.B., J.E.C., T.R.B., D.M.S., and J.L. performed and/or provided assistance with the cell-based genomic screening. T.R.P., J.P., and L.E.S. wrote the paper.

## Data and materials availability

All data associated with this study are present in the paper, the Supplementary Materials, or will be available at NCBI BioProject ID: PRJNA624650. Shared reagents are subject to a materials transfer agreement.

## SUPPLEMENTARY MATERIALS

### Materials and Methods

**Fig. S1. *ATRAID* is required for the cellular responses to nitrogen-containing bisphosphonates.**

**Fig. S2. Generation and skeletal characterization of *Atraid***^**KO**^ **mice.**

**Fig. S3. *Atraid* is required cell-autonomously for the effects of N-BP on osteoclasts in two models of osteoporosis.**

**Table S1. Results of haploid genomic screen for genes required for the response to alendronate.**

**Data file S1. Statistics for *Atraid***^**KO**^ **mice basal characterization, and statistics for bone structure, strength of ovariectomized wildtype and *Atraid***^**KO**^ **animals treated with alendronate.**

**Data file S2. Statistics for bone histomorphometry and serum bone proteins in ovariectomized and senile wildtype and *Atraid***^**KO**^ **animals treated with alendronate. Data file S3. Gene expression, sequencing, and growth phenotype data for ONJ, DTC, AFF and CRISPRi and CRISPRa studies.**

## MATERIALS AND METHODS

### Materials

Reagents were obtained from the following sources: antibodies to Rap1a (SC-1482, 1:100) from Santa Cruz Biotechnology; anti-HDJ-2/DNAJ Ab-1, Clone: KA2A5.6 (cat.# MS-225-P0, 1:2000), anti-V5 (cat.# R960-25, 1:1000), and GAPDH (14C10) Rabbit mAb (cat.# 2118S, 1:1000) from Fisher Scientific; rabbit polyclonal and monoclonal antibodies to PARP (cat.# 9532) from Cell Signaling Technology; FuGENE 6 (cat.# E2691) and Complete Protease Cocktail (cat.# 11836170001) from Roche; alendronate (cat.# 1012780-200MG), etidronate (cat.# P5248-10MG), tiludronate (cat.# T4580-50MG), Acid Phosphatase Leukocyte (TRAP) kit (cat.# 387A-1KT), SYBR Green JumpStart Taq ReadyMix (cat.# 4309155), calcein (cat.# C0875-5G), FLAG M2 affinity agarose beads (Cat.# M2220), and anti-FLAG antibody (F3165), 3X FLAG peptide (F4799), ß-glycerophosphate (cat.# 50020), ascorbic acid (cat.# A5960), RANKL (cat.# R0525), M-CSF (cat.# M9170), dexamethasone (cat.# D4902), vitamin D3 (cat.# D1530), and alizarin red (cat.# A3882-1G) from Sigma-Aldrich. Zoledronic acid (zoledronate, cat.# M 1875) from Moravek; IMDM Glutamax, α-MEM, RPMI, SuperScript II Reverse Transcriptase, Platinum Pfx, Platinum Taq DNA Polymerase and inactivated fetal calf serum (IFS) from Invitrogen; NEBNext Ultra II Q5 Master Mix (cat.# M0544L), Sal I-HF (cat.# R3138L), Not I-HF (cat.# R3189L), BstXI (cat.# R0113L), Blp I (cat.# R0585L) from New England Biolabs; NucleoSpin Blood DNA isolation kit (cat.# 740950.50) from Macherey Nagel.

### Cell lines and cell culture

KBM7 cell lines were cultured in IMDM with 10% FBS. K562 cell lines were cultured in Iscove’s Modified Dulbecco’s Medium (IMDM) or Roswell Park Memorial Institute (RPMI) medium with 10 mM Glutamine with 10% FBS. HEK-293T cells were cultured in Dulbecco’s Modified Eagle’s Medium (DMEM) with 10% FBS. RAW264.7 and primary mouse cells were cultured as described below.

### Haploid genetic screening

The genetic selection with alendronate was performed on 100 million mutagenized KBM7 cells (*53*). Cells were exposed to 165 µM alendronate and allowed to recover for 4 weeks before harvesting the surviving cells and isolating genomic DNA. Analysis of the screen was performed essentially as described previously (*53*). In brief, the sequences flanking retroviral insertion sites were determined using inverse PCR followed by Illumina sequencing. After mapping these sequences to the human genome, we counted the number of inactivating mutations (mutations in the sense orientation or present in exon) per individual Refseq-annotated gene as well as the total number of inactivating insertions for all Refseq-annotated genes. For each gene, statistical enrichment was calculated by comparing how often that gene was mutated in the drug-treated population with how often the gene carries an insertion in the untreated control dataset. For each gene, the *P*-value (corrected for false discovery rate) was calculated using the one-sided Fisher exact test (table S1).

### CRISPRi/a genetic screening

We sought to improve the gene coverage in the zoledronate CRISPRi screen we previously reported because we observed it was relatively under-sampled compared to other CRISPRi/a screens we and others had performed (*12, 54*). We isolated genomic DNA from biological replicate samples from our zoledronate screen and adapted the Weissman lab library preparation protocol (*55*) with the following two changes to achieve greater sgRNA coverage: i) we increased the input genomic DNA from 0.5 µg to 10 µg of per PCR reaction; and ii) we switched from Phusion to NEBNext Ultra II Q5 PCR enzyme. Preparing the library with these modifications didn’t change the top hits but it resulted in greater number of sgRNAs for each gene being detected and more robust overlap of the gene scores with our other CRISPRi/a screens with N-BPs. Data analysis including statistical calculations for the screens using the v2 CRISPRi/a libraries were performed as described in detail in our previous work by Zhou et al. (*12*).

### cDNA manipulations and mutagenesis

The cDNAs for *ATRAID* were cloned from a human R4 cell line cDNA library. For expression studies, all cDNAs were amplified by PCR and the products were subcloned into the Sal 1 and Not 1 sites of pRK5 or pMSCV (*56*). The controls, *metap2* and *tubulin*, were previously described (*57*). All constructs were verified by DNA sequencing. To generate the 293T *ATRAID_KO* expressing *ATRAID-FLAG*, using Gibson assembly we cloned the cDNA encoding *ATRAID* with a C-terminal Flag tag into a homologous recombination vector targeting the *AAVS1* locus, with a GGGGSGGGGS flexible linker (sequence: GGT GGA GGG GGA AGT GGC GGA GGA GGT TCA) added between the CDS and the Flag tag. We transfected this vector along with pX330 expressing an sgRNA targeting the *AAVS1* locus. To generate the *ATRAID* low expressing cell line (293T *ATRAID_KO* + *ATRAID*_*low expr*_), we used the same procedure, but cloned the cDNA encoding *ATRAID* with a C-terminal V5 tag driven by a PGK promoter. Low expression was verified with multiple *ATRAID* expression primers, described below. Mutagenesis to engineer *ATRAID* patient variants was performed with a QuikChange site-directed mutagenesis kit (Agilent) following the manufacturer’s protocol. The viral vector, pLenti PGK hygromycin *ATRAID*-V5, used for mutagenesis was previously described (*12*) and is based on Addgene clone ID: 19066.

### sgRNA manipulations

Genome editing experiments were designed based on an established protocol (*58*). In brief, the sgRNAs for *ATRAID* were cloned using Golden Gate assembly into pSpCas9-BB-2A-GFP (PX438), a kind gift from Feng Zhang (Addgene #48138). All constructs were verified by PCR and DNA sequencing. For the human *ATRAID* locus, one sgRNA targeting exon 3 and another targeting exon 5 were used to act simultaneously and remove part of exon 3, the entire exon 4 and part of exon 5.

#### Human *<u>ATRAID</u>*

exon3 sgRNA_1: GCCTGATGAAAGTTTGGACC

exon3 sgRNA_2: CCCTGGTCCAAACTTTCATC

exon5 sgRNA_1: GTCCTGGAGGAATTAATGCC

exon5 sgRNA_2: GTCCTGGAGGAATTAATGCC

#### Mouse *<u>Atraid</u>*

GGATACATCGAAGCTAATGC

For generating *Atraid* knockouts in RAW 264.7 cells, cells were transfected with PX438 carrying the above sgRNA, and clones were generated by single cell sorting. Mutation was confirmed by PCR and sequencing.

### Cell viability assays

Wild-type or mutant cells were seeded at 20,000 cells per well in a 96-well tissue culture plate and treated with indicated concentrations of compound or left untreated. 48 or 72 hours after treatment the cell viability was measured using a Cell-titer Glo colorimetric assay (Promega) according to manufacturer’s protocol. Viability is plotted as percentage viability compared to untreated control.

For assays of apoptosis, cells were plated in 6-well dishes and exposed to alendronate for 48 hours. AnnexinV positive cells were quantified using flow cytometry with the Dead Cell Apoptosis Kit with AnnexinV Alexa Fluor (488) and PI (Thermo Fisher Scientific #V13241) following the manufacturer’s instructions.

### Protein analysis

All cells unless otherwise stated were rinsed twice with ice-cold PBS and lysed with Triton-X 100 or NP-40 containing lysis buffer [40 mM HEPES (pH 7.4), 2 mM EDTA, 150 mM NaCl, 50 mM NaF, 1% Triton-X 100 or 1% NP-40, and one tablet of EDTA-free protease inhibitors (Roche) per 25 ml or Halt protease-phosphatase inhibitor (#78442 ThermoFisher Scientific)]. Lysate is incubated at 4 centigrade for 15-30min with constant inversion. The soluble fractions of cell lysates were isolated by centrifugation at 13,000 rpm for 10 min in a microcentrifuge. Lysate protein concentrations were normalized by Bradford assay (Bio-Rad). Proteins were then denatured by the addition of sample buffer and by boiling for 5 minutes, resolved using 4%-20% or 6% (for HDJ-2) SDS-PAGE (Invitrogen), and analyzed by immunoblotting for the indicated proteins. Immunoblotting was performed as follows: nitrocellulose membranes were blocked at room temperature (RT) with 5% non-fat milk for 45min. Membranes were then incubated overnight at 4°C with desired primary antibodies dissolved in 5% milk. Membranes were then washed 3 times in TBST, each wash lasting 5min. Membranes were then incubated at RT with desired secondary antibodies at 1:2000 in 5% milk for 45 minutes. HRP-conjugated or fluorescent secondary antibodies (Santa Cruz Biotechnology or Thermo Fisher, respectively) were used for detection. Membranes were then washed 3 times in TBST, each wash lasting 5min. Signal from membranes using HRP-conjugated secondary antibodies were captured using a camera and those using fluorescent secondary antibodies were imaged on a GE Typhoon Trio imager. The small GTPase Rap1A protein prenylation can be detected by immunoblot analysis using an antibody that specifically binds to its unprenylated form (*59, 60*) (Santa Cruz, SC-1482).

### RAW 264.7 differentiation

RAW 264.7 cells were maintained in DMEM + 10% FBS + 1X penicillin/streptomycin. Differentiation of RAW cells to osteoclasts was achieved following the protocol of Collin-Osdoby *et al.* (*35*) where cells were treated with 35ng/ml RANKL (R&D Systems) for 6 days. For experiments with alendronate, the drug was added at the indicated concentrations 48 hours prior to collection.

### Primary bone marrow isolation and osteoclast differentiation

Primary bone marrow was isolated from the femurs, tibiae, and spine of wildtype and *Atraid*^KO^ mice, enriched to macrophages and differentiated to osteoclasts following the protocol of Tevlin *et al.* (*34*). Where indicated, cells were treated with the indicated concentrations of alendronate 48 hours before collection. Note: alendronate responsiveness can differ between different osteoclast cell contexts (*61*). In our hands, isolated primary osteoclasts were more sensitive to alendronate than the RAW 264.7 cell line. Therefore, we used higher doses of alendronate with RAW 264.7 cells vs. isolated primary cells (30 µM and 80 µM vs. 10 µM and 30 µM).

### Co-culture of primary osteoblasts and osteoclasts

Murine bone marrow macrophages and bone marrow stromal cells were isolated from the long bones of wild-type and *Atraid*-deficient mice. Murine bone marrow macrophages were differentiated to osteoclasts using M-CSF and RANKL in α-MEM + 10% FBS. Murine bone marrow stromal cells were differentiated to osteoblasts using ß-glycerophosphate and ascorbic acid in α-MEM + 20% FBS. 200,000 osteoblasts and 300,000 osteoclasts were co-cultured in six-well dishes in α-MEM + 10% FBS 10 nM 1,25(OH)_2_ vitamin D3, and 100 nM dexamethasone and treated with the indicated concentrations of alendronate for 96 hours followed by tartrate acid phosphatase assay following the manufacturer’s protocol. Images of the 6-well dishes were captured by camera and the total TRAP staining on these images was quantified using ImageJ software using the Analyze->Analyze Particle workflow.

### Gene expression analysis

Total RNA was isolated and reverse-transcription was performed from cells or tissues in the indicated conditions. The resulting cDNA was diluted in Dnase-free water (1:20) followed by quantification by real-time PCR. mRNA transcripts were measured using Applied Biosystems 7900HT Sequence Detection System v2.3 software. All Data are expressed as the ratio between the expression of target gene to the housekeeping genes RPLP0 and/or GAPDH. Each treated sample was normalized to controls in the same cell type.

Human Primer sequences (for clarity, denoted with prefix: “h” for human):

hATRAID exon1-F’ – GGATGGAGGGGCCCGAGTTTCTG

hATRAID exon2-R’ – CCCAAGATGGTGCCCTTCTGATTC

hATRAID exon6-F’ – CCATGGATACAAGTGTATGCGCC

hATRAID exon 7-R’ – TCATGAAGTCTTGGCTTTTCGGC

hRPLP0-F’ – CAGATTGGCTACCCAACTGTT

hRPLP0-R’ – GGAAGGTGTAATCCGTCTCCAC

hTBP-F’ – GAGCCAAGAGTGAAGAACAGTC

hTBP-R’ – GCTCCCCACCATATTCTGAATCT

Mouse Primer sequences (for clarity, denoted with prefix: “m” for mouse):

m*Atraid* exon3-F’ – GATCTTCAGAACTGTTCCCTGAAG

m*Atraid* exon4-R’ – GCTGAGTAAACCCACGGAAGGTG

m*Atraid* exon5-F’ – CTTCTTTCAAGGACAAGCAGATTTG

m*Atraid* exon 7-R’ – GAATCCCAAAGAACATAAGCAGTG

m*Actb*-F’ – TGTCGAGTCGCGTCCA

m*Actb*-R’ – ATGCCGGAGCCGTTGTC

m*Rplp0*-F’ – TGCTCGACATCACAGAGCAG

m*Rplp0*-R’ – ACGCGCTTGTACCCATTGAT

m*Tnfrsf11a (RANK)*-F’ – GCAGCTCAACAAGGATACGG

m*Tnfrsf11a (RANK)-*R’ – TAGCTTTCCAAGGAGGGTGC

m*Acp5 (TRAP)*-F’ – AAGAGATCGCCAGAACCGTG

m*Acp5 (TRAP)*-R’ – CGTCCTCAAAGGTCTCCTGG

m*Ctsk*-F’ – CCTTCCAATACGTGCAGCAG

m*Ctsk*-R’ – CATTTAGCTGCCTTTGCCGT

### *Generation and genotyping of* Atraid *KO mice*

Chimeric mice were obtained by microinjection of the correctly targeted *Atraid* EUCOMM ES clones (HEPD0577_2_D01 and HEPD0577_2_E01) (*62, 63*) into BALB/C blastocysts and crossed with C57BL/6 mice to obtain offspring with germline transmission. Heterozygous mice for the floxed *Atraid* allele (*Atraid* ^loxP/+^), were crossed to C57BL/6 mice expressing the Cre-recombinase transgene from the full-body CMV promoter (*64*). Mice analyzed in this study were 100% C57BL/6.

All experiments involving mice were performed with protocols approved by the Harvard and Washington University Animal Studies Committees. We confirm that we have complied with all relevant ethical regulations. All mice were housed under a 12 hour light cycle and fed standard chow diet ad libitum.

PCR genotyping of all WT and *Atraid* deficient mice were performed with primers that detect the following:

1. This generates a 140bp product and indicates the presence of the transgene. Transgene (92 upstream) F’ CAGCCATATCACATCTGTAGAG Transgene (92 upstream) R’ GAGTTTGGACAAACCACAACTAG
2. This indicates recombination and is detectable in mice also expressing CRE Del F’ CTGCATTCTAGTTGTGGTTTGTCC Del R’ CAGGAGGTAGTGCAAGCCTTTG
3. Wild-type *Atraid* primers spanning *Atraid* exons 3 and 4. This PCR product is not detectable in homozygous null animals. Exon ¾ F’ CAGAACTGTTCCCTGAAGGATCCTGGTC Exon ¾ R’ GTACACACTGTTAGCGCTCTGTTTGC
4. These generic CRE primers give a ∼100bp product indicates the presence of the CRE transgene. CRE F’ GCG GTC TGG CAG TAA AAA CTA TC CRE R’ GTG AAA CAG CAT TGC TGT CAC TT

### Serum ELISA assays

Cardiac puncture blood of mice of the indicated ages was obtained and centrifuged at low speed at 4°C for 15 minutes, and serum was isolated. Gla-Osteocalcin (Mouse Gla-Osteocalcin High Sensitive EIA Kit from Clontech, cat.# MK127) and C-terminus cross-linked telopeptides of type I collagen CTX-I (RatLaps EIA Kit from Immunodiagnosticsystems Inc., AC-06F1) were quantified following the manufacturer instructions.

### Animal procedures

Age and sex matched mice were randomly assigned to treatment groups within each genotype. For the basal characterization of WT and *ATRAID* deficient mice and for the OVX experiments the mice were litter-matched. For the senile osteoporosis experiments the animals were originally derived from the same litter but were bred as cohorts. All animal experiments were replicated two to four times spanning independent days to ensure reproducibility. No outlier animals were excluded from any downstream analysis.

Ovariectomy or sham operations were performed on 3.5-month-old females as detailed previously (*65*). Briefly, the ovaries were exposed through an abdominal approach and either resected after clipping the blood vessels or left in place (sham operation). The muscle and skin of the abdomen were sutured. Mice were given an intraperitoneal injection of buprenex immediately after surgery and then every twelve hours for 48 hours post-surgery. Immediately preceding OVX, vehicle (phosphate buffered saline) or 100 μg/kg alendronate (both provided by Sigma) was injected intra-peritoneally every week for 4 weeks. These doses were chosen based on the anti-resorptive activity of alendronate in different species (*66, 67*). Mice utilized for the senile model of osteoporosis were given the same regime of alendronate for 4 weeks starting at 18 months old. Power calculations and cohort sizes for the senile model experiments were based on the ovariectomy (OVX) model experiment statistical calculations. We acknowledge larger N’s would be preferable for the senile model as it is in most experiments. However, it was not feasible within a reasonable time frame to obtain more litter-, sex-, and genotype-matched animals for these studies, especially to reach the age we used, 18 months old to meet the minimum criteria for the mice to be considered “senile”. Also, because the referenced experiment was a 2^nd^ supporting model on the more widely used OVX model we felt confident proceeding with the experiment despite it being relatively modestly powered.

### Bone microstructure

A high-resolution desktop micro-tomographic imaging system (µCT40, Scanco Medical AG) was used to assess cortical and trabecular bone microarchitecture and morphology in the femoral mid-diaphysis and distal metaphysis, respectively. Scans were acquired using a 10 µm^3^ isotropic voxel size, 70 kVP peak x-ray tube potential, 114 mAs x-ray intensity, 200 ms integration time, and were subjected to Gaussian filtration and segmentation. Regions of interest (ROIs) were selected 50 slices above and below the femoral longitudinal midpoint or 100 slices above the growth plate of the distal femur to evaluate the cortical and trabecular compartment, respectively. Image acquisition and analysis adhered to the JBMR guidelines for the use of µCT for the assessment of bone microarchitecture in rodents (*24*). To judge the effect sizes in our µCT experiments, it is notable that both OVX and alendronate influenced µCT parameters in wild-type mice of a magnitude in our hands (∼10-20%) that is highly consistent with what has been previously shown in the strain we used, C57BL/6 (*32, 68*).

### Bone biomechanics

Mechanical testing in a 3-point bending to failure was conducted on femora after µCT. Briefly, hydrated femora were stabilized over supports 7mm apart and a loading force was applied in the anteroposterior direction midway between the supports (Instron). Test curves were analyzed to determine ultimate force to failure and stiffness as described previously (*25, 69*).

### Bone histomorphometry

To label mineralizing fronts, mice were injected intraperitoneally with calcein (15 mg/kg i.p., Sigma-Aldrich) and alizarin red (30 mg/kg; Sigma) were intraperitoneally injected 7 and 2 days, respectively, before euthanasia. Bone was fixed in 10% (vol/vol) neutral buffered formalin for 24 hours, processed through a series of graded alcohols, and decalcified. Decalcified vertebrae or femurs were embedded in paraffin and 2-4 hours, processed through a series of graded alcohols, and Tartrate resistant acid phosphatase (TRAP) stain was performed. Undecalcified femora were embedded in methyl methacrylate and the whole bone were cut and stained for TRAP or analyzed for calcein and alizarin red fluorescence. Quantitative histomorphometry was performed using a commercial software (OSTEO II, Bioquant), and standard parameters of bone remodeling were determined as detailed elsewhere (*70*).

### Transcriptional profiling analysis

The multiple myeloma, osteonecrosis of the jaw (ONJ) microarray gene expression was performed as previously described (*36*) using the Affymetrix U133 Plus 2.0 array platform (Affymetrix) on total RNA isolated from peripheral blood mononuclear cells (GEO accession #: GSE7116). 21 multiple myeloma patients with a history of N-BP use were included in the study. 11 patients (52.4%) reported to have ONJ. The breast cancer bone marrow micrometastases (also known as DTC) microarray gene expression data was generated as previously described (*37*) and also used the Affymetrix U133 Plus 2.0 array platform. DTC profiling was performed on tumor biopsies of 81 patients (GEO accession #: GSE71258). 54 breast cancer patients treated with zometa (zoledronate) were included in the study. N-BPs directly inhibit tumor growth and angiogenesis (*71, 72*). 14 patients (25.9%) eventually died and 40 patients (74.1%) survived. Her-2 negative patients were divided into two categories following randomization to the zoledronate arm: those that who had DTC reoccurrences or lived less than 1000 days and those who lived at least 2500 days.

Quantile normalization was used for all differential expression analysis, and all the normalization procedures were performed using function normalizeQuantiles in the R Bioconductor package limma (*73*). Gene expression data was filtered using function filterfun in the R Bioconductor package genefilter (*74*). Probes with expression values over 5 in less than 25% of the samples were removed. Comparison between groups were estimated using an empirical Bayes method (*75*), and variances across all genes were used to estimate the variance of each gene. Raw *P*-values were calculated from a moderated t-test, and false discovery rate (FDR) adjusted *P*-values were obtained based on Benjamini and Hochberg’s methods for multiple testing. Log_2_ fold changes between the experimental conditions were calculated for each gene as well.

Affy probe IDs were transformed into gene symbols based on the R Bioconductor package, hgu133plus2.db (*76*). In Fig. 4A, differentially regulated genes for the -/+ONJ patients were identified by having adjusted *P*-values smaller than 0.05, while potential differentially-regulated genes for the <1000 vs. >2500 days with breast cancer patients with disseminated tumor cells (DTC) were identified by having raw *P*-value smaller than 0.05. For the ONJ dataset, 1992 genes were significant. For the DTC dataset, 1854 genes were significant.

### Exome sequencing analysis

Exome sequencing data was generated from blood leukocyte DNA for 27 bisphosphonate treated osteoporosis cases with atypical femoral fractures (AFF), 11 bisphosphonate treated osteoporosis cases without atypical femoral fractures, and 8 bisphosphonate multiple myeloma or breast cancer cases with osteonecrosis of the jaw (ONJ). Exome capture was performed using Agilent All-exome capture kits. Sequencing was performed using paired-end Illumina sequencing. Analysis of exome sequencing data was performed in-house using our previously described methods (*77, 78*). Briefly, FASTQ formatted sequences were aligned to the hg37 human reference sequence (NCBI GRCh37) using BWA (*79*). Mapped reads were filtered to remove duplicate reads with the same paired start sites. The Binary sequence Alignment/Map (BAM) formatted alignments were then processed using the Genome Analysis Toolkit (GATK) Haplotype Caller (*80, 81*) and genotypes jointly called together with all in-house control exome sequenced individuals. Variants were filtered for read-depth (>8x), genotype quality (GQ>20), and GATK-calculated variant quality score recalibration (VQSR). Allele frequencies were annotated using the gnomAD database (*82*). Variant positions were processed excluding those with call rates < 0.95 or Hardy-Weinberg equilibrium *P*-values < 10^−5^. Variants were annotated using Seattleseq: http://snp.gs.washington.edu/SeattleSeqAnnotation151/. Genes with multiple variants with non-Finnish European (NFE) allele frequencies less than 0.05 only in cases were considered. From dbSNP, https://www.ncbi.nlm.nih.gov/SNP/: For the *ATRAID* gene, the D5G variant is found on chromosome 2 position: 27212382 – rs1275533. The G32R variant is found on chromosome 2 position: 27212297 – rs11556163.

The ethnic breakdown of the AFF cases is: 25/27 European American (EA), 3/27 Asian American (AA), 0/27 African American. The ethnic breakdown of the controls is 8/11 European American and 3/11 unknown. For *ATRAID*, the D5G variant is present in 2 out of 27 AFF patients. Though this allele wasn’t detected in the 11 control samples, it is present in a population of European Americans (EA) that is representative of the study population at a prevalence of 0.0139 and in Asian Americans (AA) at a prevalence of 0.002. Therefore, the D5G allele is (2/27) / (0.0139 * 25/27 + 0.002 * 3/27) = 5.66X enriched in cases compared to a representative population to that of the cases. For the G32R *ATRAID* allele the enrichment is: (2/27) / (0.0149 * 25/27 + 0.002 * 3/27) = 5.28. Because we have incomplete information on the ONJ cases, we used the EA frequency for our enrichment calculations: the D5G allele is (1/8) / (0.0139 * 8/8) = 7.19X enriched in cases compared to a representative population. For the G32R allele the enrichment is: (1/8) / (0.0149 * 8/8) = 6.71X. A simple binomial test, binom.test(), was used to calculate the *P*-value for each *ATRAID* variant in AFF and ONJ cases. For the D5G allele, *P*-value = 0.01262 – binom.test(3, 35, 0.0139). For the G32R, *P*-value = 0.01518 – binom.test(3, 35, 0.0149).

### Functional validation of ATRAID patient alleles

For testing the effects of reduced *ATRAID* expression, *ATRAID* deficient HEK-293T cells were stably infected with sub-endogenous expression of variant “v3” of wild-type *ATRAID*. These cells were then compared with wild-type HEK-293T and the parent *ATRAID* deficient line in cell viability assays. For the rs1275533 variant, the amino acid change, D5G, corresponds to the v3 ATRAID amino acid sequence and the same variant is D60G on the longer, v2 *ATRAID* isoform. For the rs11556163 variant, G32R is only present on v2 *ATRAID*. Both *ATRAID* variants were found together in each AFF or ONJ case identified, which suggested these variants are linked. Thus, to recapitulate the patient genotypes, we therefore introduced both v2 and v3 wild-type or variant forms of ATRAID in *ATRAID* deficient cells. This means there were three differences between the wild-type and ATRAID D5G/G32R variant cell lines we generated – two variants on v2 ATRAID and one variant on v3 ATRAID.

## SUPPLEMENTARY FIGURES

**Fig. S1.**
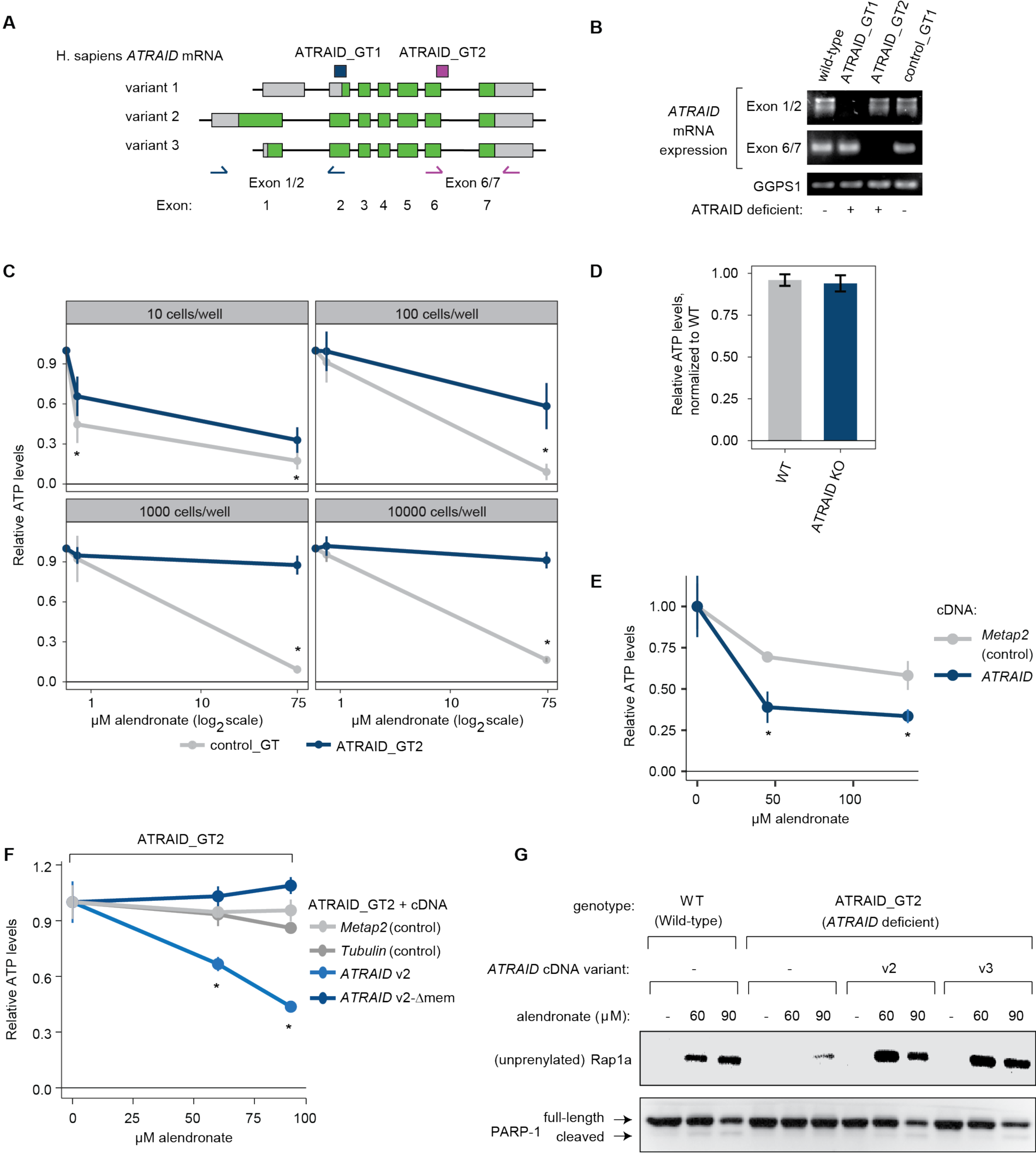
*ATRAID* is required for the cellular responses to nitrogen-containing bisphosphonates. **(A)** Schematic of the exon structure of the three human *ATRAID* mRNA variants. The coding sequence for each variant is in green. Non-coding portions of each exon are in grey. The translated regions of variant 1 and variant 3 are shorter than variant 2 due to internal translation initiation sites. The location of the primer sets used to identify each *ATRAID* gene trap (GT) are indicated (in blue for ATRAID_GT1; in red for ATRAID_GT2). **(B)** mRNA analysis of *ATRAID* and *GGPS1* expression in clones that contain independent gene-trap insertions in their respective loci. Wild-type KBM7 cells were compared with mutant alleles (labeled as GT) and *GGPS1* was used as a loading control. **(C)** The growth inhibitory effects of alendronate on WT and *ATRAID* deficient cells over a wide range of concentrations and cell numbers. Viability was determined by measuring cellular ATP and expressed as a ratio of that compared with untreated cells. All measurements were performed in quadruplicate (biological replicates). \**P* < 0.05, student’s *t*-test. **(D)** Cell growth rate of untreated WT and *ATRAID*-deficient cells. WT and *ATRAID*-deficient HEK-293T cells were plated at the same cell density (10,000 cells/per well), allowed to grow for 48 hours, and ATP was measured. Viability measurements were performed in quadruplicate (biological replicates). Error bars reflect standard deviation. **(E)** The growth inhibitory effects of alendronate on *ATRAID* overexpressing cells. HEK-293T cells were transfected with either Metap2 (control), or *ATRAID*-cDNA expressing vectors and the indicated doses of alendronate for 72 hours. Viability measurements were performed in quadruplicate (biological replicates). \**P* < 0.05, student’s *t*-test. **(F)** The growth inhibitory effects of alendronate on membrane-targeted vs non-targeted *ATRAID* expressing cells. Cells deficient in *ATRAID* (ATRAID_GT2) were transformed to express exogenous Metap2 (control), tubulin (control), ATRAID variant 2 (_v2), or ATRAID variant 2 lacking the transmembrane domain (Δmem_v2). Cells were treated with alendronate at the indicated dose for 72 hours. Cell viability was determined as in C. \**P* < 0.05, student’s *t*-test. (n=6) (3 biological replicates, 3 technical replicates). **(G)** The effects of alendronate and *ATRAID* deficiency on prenylation in an additional cell type, KBM7. Wild-type control and *ATRAID* deficient KBM7 cells exogenously expressing or not expressing *ATRAID* cDNA were treated with the indicated dose of alendronate for 24 hours then lysed and analyzed by immunoblotting for the indicated proteins. Equal amounts of protein were loaded in each lane. This experiment was repeated three times (biological replicates) and was consistent all three times.

**Fig. S2.**
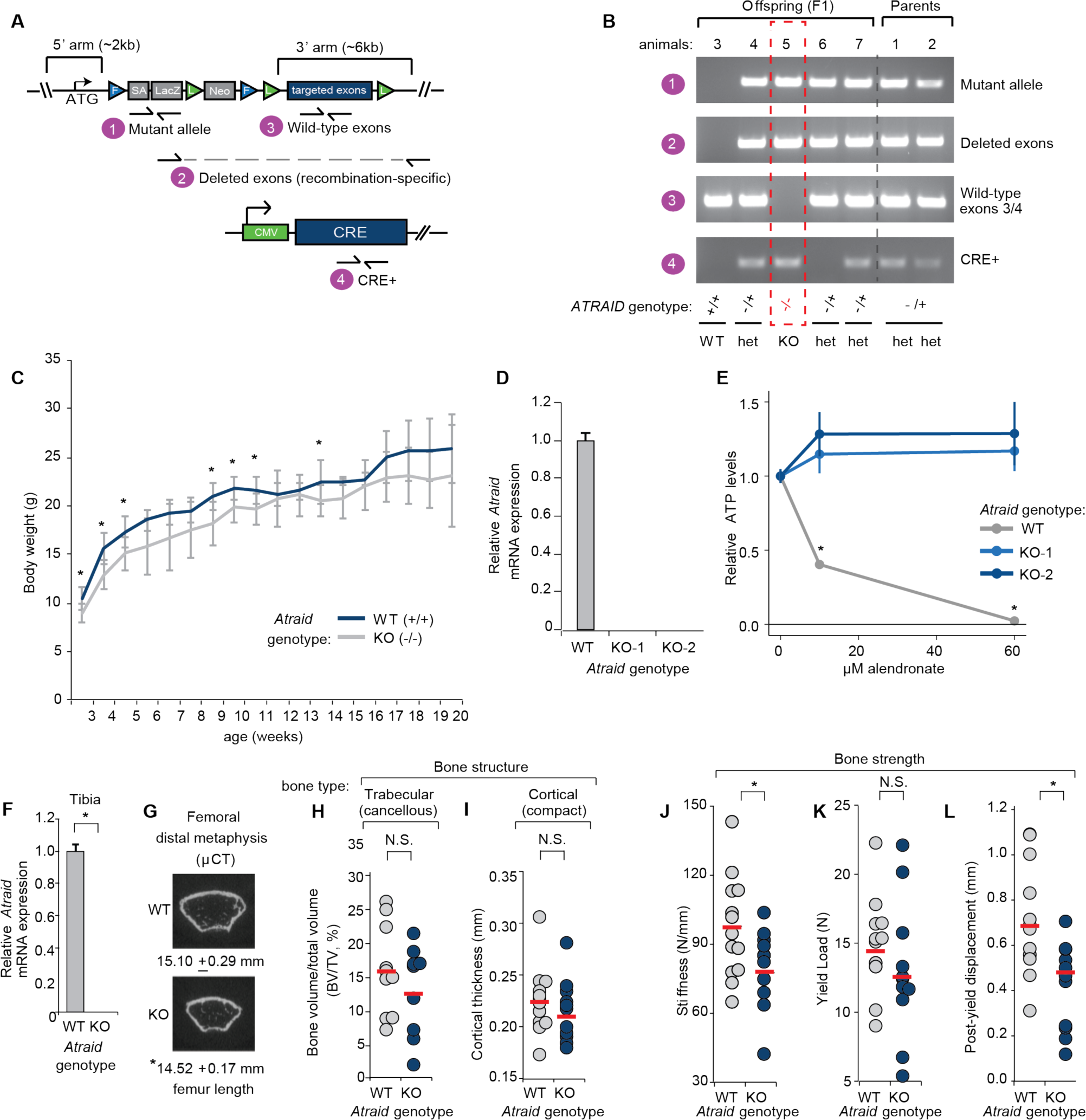
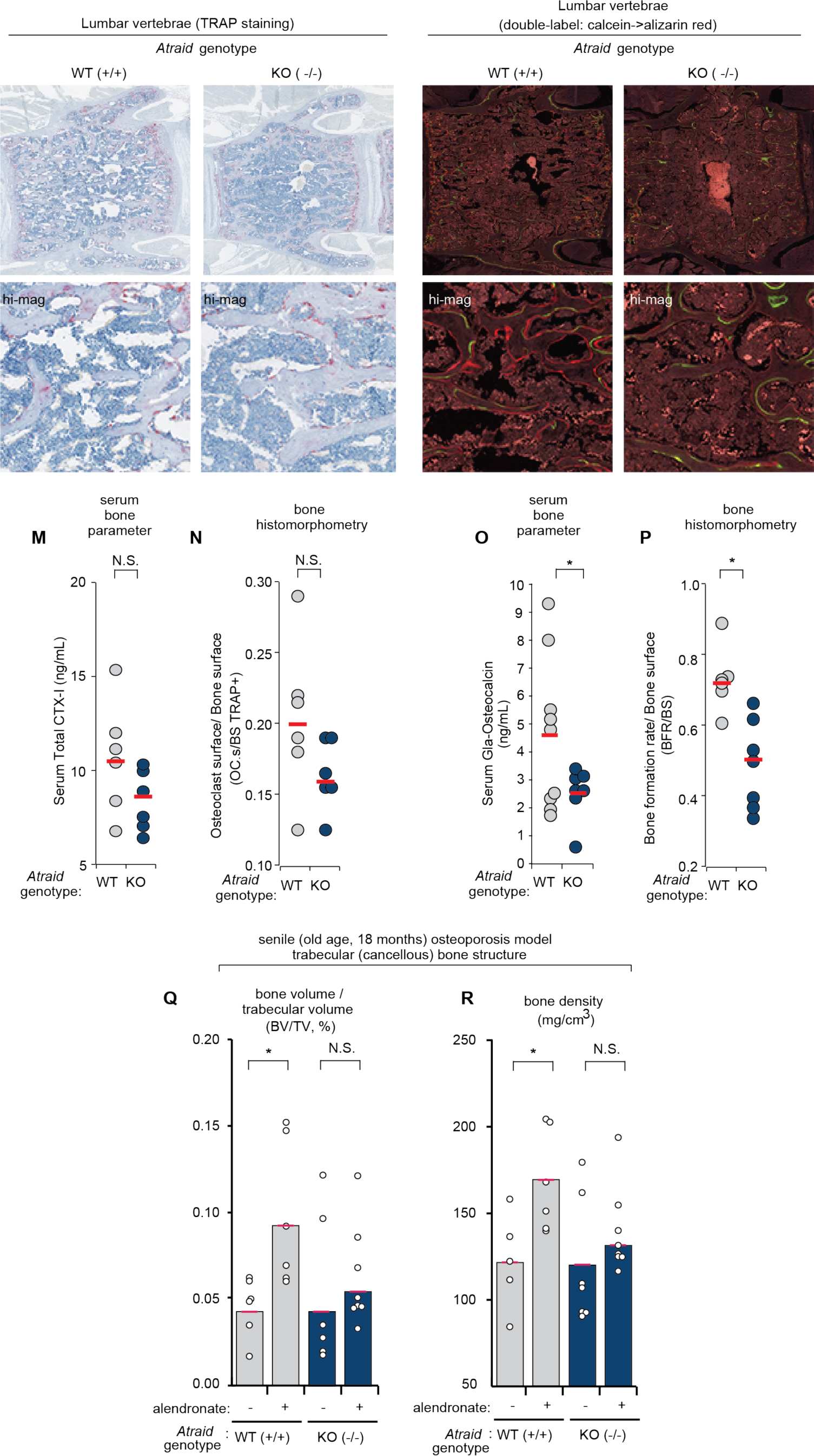
Generation and skeletal characterization of *Atraid*^KO^ mice. **(A)** Schematic of *ATRAID* targeted gene locus and genotyping strategy. *Atraid* exon 3-exon 5 are targeted, flanked by LoxP sites. **(B)** Assessment of deletion of protein-coding genomic DNA at the *Atraid* gene locus. WT, wildtype at the *Atraid* locus; het, heterozygous; KO, homozygous deletion. **(C)** Body weights of *Atraid* KO (-/-) and wild-type, WT (+/+) mice from 3 to 20 weeks of age. N=6-13 for wild-type, N=7-14 for *Atraid*^KO^ mice. \**P* < 0.05, student’s *t-*test. **(D)** *Atraid* mRNA expression in WT and *Atraid*^KO^ mouse tail fibroblast cells were analyzed by RT-qPCR and normalized to *Rplp0* expression. Error bars indicate standard deviation for n=4 (biological replicates). * indicates *P*<0.05, student’s *t-*test. **(E)** The growth inhibitory effects of alendronate of cells from the tails of WT and *Atraid*^KO^ mice. All cells were treated with the indicated concentration of alendronate for 72 hours. Cell viability was determined by measuring cellular ATP and is expressed as a ratio of that compared with untreated cells. All measurements were performed in quadruplicate (biological replicates). \**P* < 0.05, student’s *t-*test. **(F)** *Atraid* mRNA in tibia of WT and *Atraid*^KO^ mice was analyzed by RT-qPCR and normalized to *Rplp0* mRNA expression. Error bars indicate standard deviation for n=4 (biological replicates). \**P* < 0.05, unpaired t-test. **(G)** Representative traverse µCT images at the femoral distal metaphysis from wild-type and *Atraid*^KO^ mice. Femur lengths in millimeters (mm) were based on µCT measurement. Error measurements are standard deviation for n=5-6 mice. * indicates *P*<0.05, student’s *t-*test. **(H, I)** Bone microstructure in *Atraid*^KO^ and WT mice. Femur trabeculae (H) and cortical (I) regions were analyzed by µCT. Each circle represents an individual animal. Circles offset to the right represent unique animals with similar values to those of another animal (offset for visual clarity). N=8-10 (2-month-old) mice per group. n.s. indicates not significant, student’s *t*-test. **(J-L)** Bone strength in *Atraid*^KO^ and WT mice. Stiffness (J), yield load (K), and post-yield displacement (L) were analyzed by three-point bending test. Each circle represents an individual animal. Circles offset to the right represent unique animals with similar values to those of another animal (offset for visual clarity). N=8-10 mice (2-month-old) per group. \**P* < 0.05, n.s. indicates not significant, student’s *t-*test. **(M, N)** Markers of osteoclast function in *Atraid*^KO^ and WT mice. (M) Serum C-terminal telopeptides of type I collagen (CTX-I) were measured in serum obtained from 3-month-old males using ELISA. Each circle represents an individual animal. Circles offset to the right represent unique animals with similar values to those of another animal (offset for visual clarity). N=6 mice per group. n.s. indicates not significant, student’s *t-*test. (N) Osteoclast surface to bone surface ratio (Oc.S/BS) was determined by Tartrate Acid Phosphatase (TRAP)-assay reactivity. Each circle represents an individual animal. Circles offset to the right represent unique animals with similar values to those of another animal (offset for visual clarity). N=6 mice per group. n.s. indicates not significant, student’s *t-*test. **(O, P)** Markers of osteoblast function in in *Atraid*^KO^ and WT mice. (O) Gla-Osteocalcin was measured in serum obtained from 3-month-old males using ELISA. Each circle represents an individual animal. Circles offset to the right represent unique animals with similar values to those of another animal (offset for visual clarity). N=7-9 mice per group. \**P* < 0.05, student’s *t-*test. (P) Bone formation rate/bone surface (BFR/BS), determined by double labeling using calcein followed by alizarin red, was analyzed histologically. Each circle represents an individual animal. Circles offset to the right represent unique animals with similar values to those of another animal (offset for visual clarity). N=5-7 mice per group. \**P* < 0.05, n.s. indicates not significant, student’s *t-*test. **(Q, R)** Bone microstructural responses to alendronate in senile osteoporotic (18 month old) *Atraid*^KO^ and WT mice. Femur trabeculae regions from WT and *Atraid*^KO^ mice were analyzed by µCT. Each circle represents an individual animal. Circles offset to the right represent unique animals with similar values to those of another animal (offset for visual clarity). N=5-8 mice per group. \**P* < 0.05, student’s *t-*test, and red line indicates mean.

**Fig. S3.**
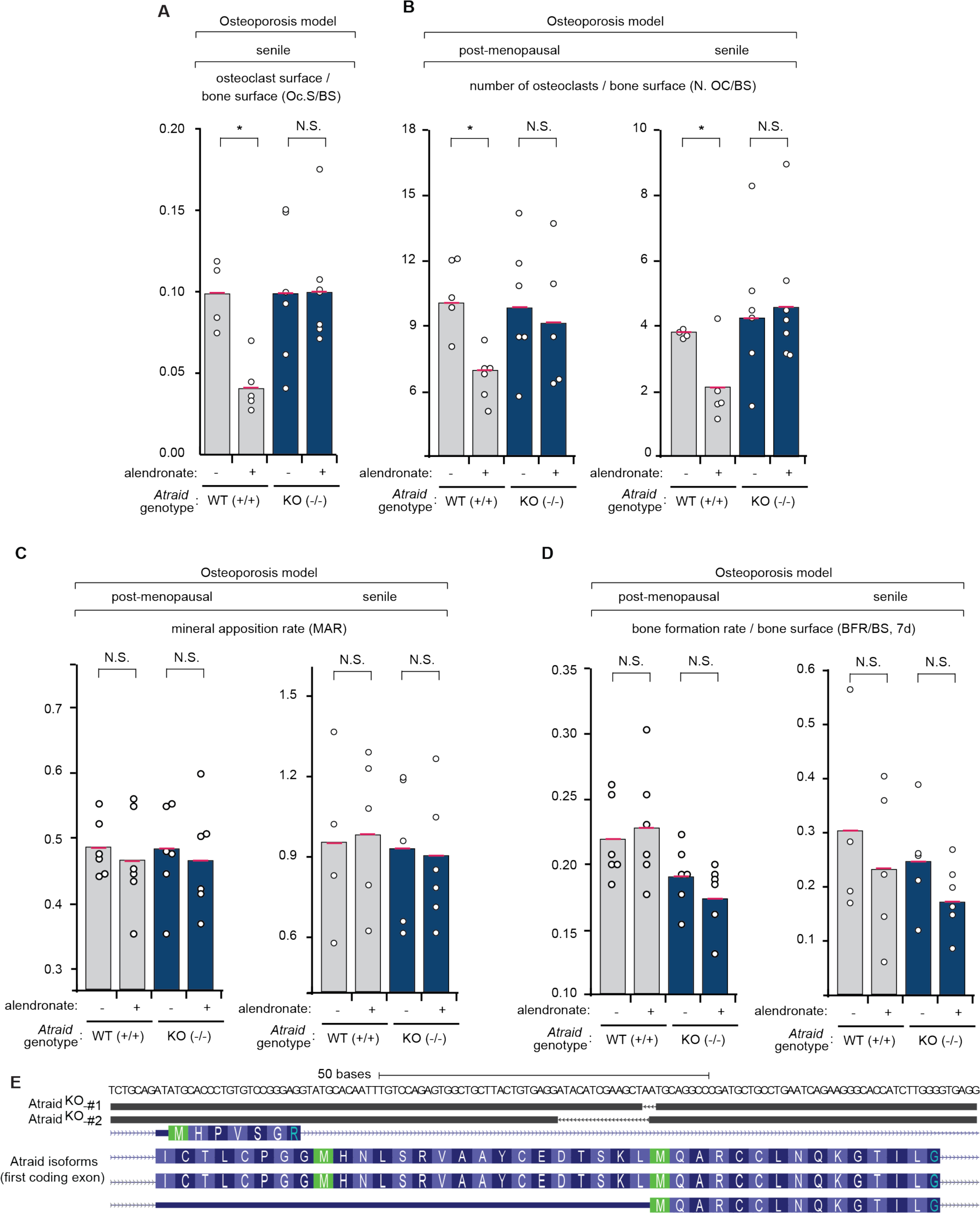
*Atraid* is required cell-autonomously for the effects of N-BP on osteoclasts in two models of osteoporosis. **(A, B)** Osteoclast surface and numbers are in WT and *Atraid*^KO^ ovariectomized or senile mice by alendronate treatment. Osteoclast surface per bone surface (Oc.S/BS) (A) and the number of osteoclasts per bone surface (N.Oc/BS) (B) were determined by Tartrate-Resistant Acid Phosphatase (TRAP)-assay reactivity. Each circle represents an individual animal. Circles offset to the right represent unique animals with similar values to those of another animal (offset for visual clarity). N=5-7 mice per group (OVX); N=4-7 mice per group (senile). \**P* < 0.05, n.s. indicates not significant, student’s *t-*test and red line indicates mean. **(C, D)** Osteoblast function after alendronate treatment in WT and *Atraid*^KO^ ovariectomized or senile mice. Mineral apposition rate (MAR) (C) and bone formation rate (BFR/BS) (D) were determined by double labeling using Calcein followed by Alizarin Red were analyzed histologically. Each circle represents an individual animal. Circles offset to the right represent unique animals with similar values to those of another animal (offset for visual clarity). N=5-7 mice per group (OVX); N=4-7 mice per group (senile). \**P* < 0.05, n.s. indicates not significant, and red line indicates mean. **(E)** Schematic of the mouse *Atraid* first coding exon, indicating the CRISPR-induced mutations present in the two *Atraid*^KO^ clones used in Fig. 3.

**Table S1. Results of haploid genomic screen for genes required for the response to alendronate.**

Tab **(A)** List of abbreviations used in this work. **(B)** Haploid genomic screen results. Enrichment of gene-trap insertions, listed by gene, and ranked by *P*-value enrichment in the alendronate treated vs. untreated populations of cells. After mapping these sequences to the human genome, we counted the number of inactivating mutations (mutations in the sense orientation or present in exon) per individual Refseq-annotated gene as well as the total number of inactivating insertions for all Refseq-annotated genes. For each gene, the *P*-value (corrected for false discovery rate) was calculated using the one-sided Fisher exact test (table S1).

Tab **(A)** Data related to Figure 2. Top rows list mean and standard deviation of: cortical thickness, cortical area, trabecular thickness, bone volume/total volume, stiffness, and yield load, from wildtype (WT) and *Atraid*^KO^ (KO) ovariectomized mice treated with vehicle (VEH, saline) or alendronate (ALN). Below rows refer to minimum, 1^st^ quartile, median, 3^rd^ quartile, and maximum of those same measurements. N=6-11 mice (3.5 month old) per group. **(B)** Data related to Figure S2, basal characterization of wildtype and *Atraid*^KO^ 2-month-old male mice. Top rows list mean and standard deviation of bone volume/total volume, cortical thickness, stiffness, yield load, post-yield displacement, serum total CTX-I, osteoclast surface/bone surface, serum Gla-osteocalcin, and bone formation rate/bone surface in wildtype and *Atraid*^KO^ mice. Below rows refer to minimum, 1^st^ quartile, median, 3^rd^ quartile, and maximum of bone volume/total volume, and bone mineral density. N=8-10 mice per group.

**Data file S2. Statistics for bone histomorphometry and serum bone proteins in ovariectomized and senile wildtype and *Atraid***^**KO**^ **animals treated with alendronate.**

Tab **(A)** Bone histomorphometry data, related to Figure 3. Top columns list mean and standard deviation of Serum total CTX-I, osteoclast surface/bone surface from wildtype (WT) and *Atraid*^KO^ (KO) ovariectomized mice treated with vehicle (VEH, saline) or alendronate (ALN). Middle rows refer to minimum, 1^st^ quartile, median, 3^rd^ quartile, and maximum of those same measurements. N=6-11 mice (3.5 month old) per group. N=8-13 mice per group. Bottom rows refer to minimum, 1^st^ quartile, median, 3^rd^ quartile, and maximum counts of # of TRAP positive osteoclasts after of co-culture experiments of wildtype primary osteoblasts with indicated genotype of primary osteoclasts, with indicated alendronate treatment (ALN). **(B)** Bone histomorphometry data, related to Figure S3. Top rows list mean and standard deviation of osteoclast surface/bone surface, number of osteoclasts/bone surface, mineral apposition rate, and bone formation rate from wildtype (WT) and *Atraid*^KO^ (KO) 18 month-old senile mice treated with vehicle (VEH, saline) or alendronate (ALN). This is followed by rows detailing the minimum, 1^st^ quartile, median, 3^rd^ quartile, and maximum of those same measurements. Bottom rows refer to number of osteoclasts/bone surface, mineral apposition rate, and bone formation rate from wildtype (WT) and *Atraid*^KO^ (KO) ovarietomized mice treated with vehicle (VEH, saline) or alendronate (ALN). N=5-7 mice per group (OVX); N=4-7 mice per group (senile).

**Data file S3. Gene expression, sequencing, and growth phenotype data for ONJ, DTC, AFF and CRISPRi and CRISPRa studies.**

All data related to Figure 4. Tab **(A)** Gene expression from bisphosphonate treated patients who did and did not suffer osteonecrosis of the jaw (ONJ) (*36*). Tab **(B)** Whole exome sequencing bisphosphonate treated cancer patients who experienced ONJ. Tab **(C).** Gene expression from bisphosphonate treated patients who did and did not suffer bone marrow-disseminated tumor cells (DTC) **(*37*).** Tab **(D)** Zoledronate CRISPRi screen results. Mann–Whitney *U* tests determined statistically significant genes. Tab **(E)** Gene-level data on the three genes, ATRAID, ATR, and ZBTB4 that scored across all genome-wide data types – gene expression from patients, CRISPRi screening in human cells, and exome sequencing of patients. Tab **(F)** Atypical femoral fracture (AFF) patient information on the patients that were exome sequenced. Tab **(G)** ONJ patient information on the patients that were exome sequenced.

## REFERENCES

1. M. J. Favus, Bisphosphonates for osteoporosis. N Engl J Med 363, 2027–2035 (2010).

2. C. J. Rosen, Clinical practice. Postmenopausal osteoporosis. N Engl J Med 353, 595–603 (2005).

3. R. Coleman, The use of bisphosphonates in cancer treatment. Ann N Y Acad Sci 1218, 3–14 (2011).

4. N. B. Watts, D. L. Diab, Long-term use of bisphosphonates in osteoporosis. J Clin Endocrinol Metab 95, 1555–1565 (2010).

5. Y. Bi, Y. Gao, D. Ehirchiou, C. Cao, T. Kikuiri, A. Le, S. Shi, L. Zhang, Bisphosphonates cause osteonecrosis of the jaw-like disease in mice. Am J Pathol 177, 280–290 (2010).

6. S. Khosla, J. A. Cauley, J. Compston, D. P. Kiel, C. Rosen, K. G. Saag, E. Shane, Addressing the Crisis in the Treatment of Osteoporosis: A Path Forward. J Bone Miner Res, (2016).

7. S. Jha, Z. Wang, N. Laucis, T. Bhattacharyya, Trends in Media Reports, Oral Bisphosphonate Prescriptions, and Hip Fractures 1996-2012: An Ecological Analysis. J Bone Miner Res 30, 2179–2187 (2015).

8. A. A. Reszka, G. A. Rodan, Nitrogen-containing bisphosphonate mechanism of action. Mini Rev Med Chem 4, 711–719 (2004).

9. V. Breuil, F. Cosman, L. Stein, W. Horbert, J. Nieves, V. Shen, R. Lindsay, D. W. Dempster, Human osteoclast formation and activity in vitro: effects of alendronate. J Bone Miner Res 13, 1721–1729 (1998).

10. Z. Zimolo, G. Wesolowski, G. A. Rodan, Acid extrusion is induced by osteoclast attachment to bone. Inhibition by alendronate and calcitonin. J Clin Invest 96, 2277–2283 (1995).

11. D. E. Hughes, K. R. Wright, H. L. Uy, A. Sasaki, T. Yoneda, G. D. Roodman, G. R. Mundy, B. F. Boyce, Bisphosphonates promote apoptosis in murine osteoclasts in vitro and in vivo. J Bone Miner Res 10, 1478–1487 (1995).

12. Z. Yu, L. E. Surface, C. Y. Park, M. A. Horlbeck, G. A. Wyant, M. Abu-Remaileh, T. R. Peterson, D. M. Sabatini, J. S. Weissman, E. K. O’Shea, Identification of a transporter complex responsible for the cytosolic entry of nitrogen-containing-bisphosphonates. Elife 7, (2018).

13. J. E. Carette, C. P. Guimaraes, M. Varadarajan, A. S. Park, I. Wuethrich, A. Godarova, M. Kotecki, B. H. Cochran, E. Spooner, H. L. Ploegh, T. R. Brummelkamp, Haploid genetic screens in human cells identify host factors used by pathogens. Science 326, 1231–1235 (2009).

14. A. J. Roelofs, K. Thompson, F. H. Ebetino, M. J. Rogers, F. P. Coxon, Bisphosphonates: molecular mechanisms of action and effects on bone cells, monocytes and macrophages. Curr Pharm Des 16, 2950–2960 (2010).

15. F. Zhu, W. Yan, Z. L. Zhao, Y. B. Chai, F. Lu, Q. Wang, W. D. Peng, A. G. Yang, C. J. Wang, Improved PCR-based subtractive hybridization strategy for cloning differentially expressed genes. Biotechniques 29, 310–313 (2000).

16. S. Bashiardes, R. Veile, M. Allen, C. A. Wise, M. Dobbs, J. A. Morcuende, L. Szappanos, J. A. Herring, A. M. Bowcock, M. Lovett, SNTG1, the gene encoding gamma1-syntrophin: a candidate gene for idiopathic scoliosis. Hum Genet 115, 81–89 (2004).

17. Y. Z. Liu, S. G. Wilson, L. Wang, X. G. Liu, Y. F. Guo, J. Li, H. Yan, P. Deloukas, N. Soranzo, U. Chinappen-Horsley, A. Cervino, F. M. Williams, D. H. Xiong, Y. P. Zhang, T. B. Jin, S. Levy, C. J. Papasian, B. M. Drees, J. J. Hamilton, R. R. Recker, T. D. Spector, H. W. Deng, Identification of PLCL1 gene for hip bone size variation in females in a genome-wide association study. PLoS One 3, e3160 (2008).

18. E. Shimizu, J. Tamasi, N. C. Partridge, Alendronate affects osteoblast functions by crosstalk through EphrinB1-EphB. J Dent Res 91, 268–274 (2012).

19. G. Yang, F. Yu, H. Fu, F. Lu, B. Huang, L. Bai, Z. Zhao, L. Yao, Z. Lu, Identification of the distinct promoters for the two transcripts of apoptosis related protein 3 and their transcriptional regulation by NFAT and NFkappaB. Mol Cell Biochem 302, 187–194 (2007).

20. http://www.sbg.bio.ic.ac.uk/phyre2/html/page.cgi?id=index.

21. R. G. Russell, Bisphosphonates: the first 40 years. Bone 49, 2–19 (2011).

22. K. Thompson, M. J. Rogers, F. P. Coxon, J. C. Crockett, Cytosolic entry of bisphosphonate drugs requires acidification of vesicles after fluid-phase endocytosis. Mol Pharmacol 69, 1624–1632 (2006).

23. W. C. Skarnes, B. Rosen, A. P. West, M. Koutsourakis, W. Bushell, V. Iyer, A. O. Mujica, M. Thomas, J. Harrow, T. Cox, D. Jackson, J. Severin, P. Biggs, J. Fu, M. Nefedov, P. J. de Jong, A. F. Stewart, A. Bradley, A conditional knockout resource for the genome-wide study of mouse gene function. Nature 474, 337–342 (2011).

24. M. L. Bouxsein, S. K. Boyd, B. A. Christiansen, R. E. Guldberg, K. J. Jepsen, R. Muller, Guidelines for assessment of bone microstructure in rodents using micro-computed tomography. J Bone Miner Res 25, 1468–1486 (2010).

25. https://cpb-us-w2.wpmucdn.com/sites.wustl.edu/dist/f/1982/files/2019/05/Understanding-3pt-Bending-outcomes.pdf.

26. C. Rosenquist, C. Fledelius, S. Christgau, B. J. Pedersen, M. Bonde, P. Qvist, C. Christiansen, Serum CrossLaps One Step ELISA. First application of monoclonal antibodies for measurement in serum of bone-related degradation products from C-terminal telopeptides of type I collagen. Clin Chem 44, 2281–2289 (1998).

27. D. W. Dempster, J. E. Compston, M. K. Drezner, F. H. Glorieux, J. A. Kanis, H. Malluche, P. J. Meunier, S. M. Ott, R. R. Recker, A. M. Parfitt, Standardized nomenclature, symbols, and units for bone histomorphometry: a 2012 update of the report of the ASBMR Histomorphometry Nomenclature Committee. J Bone Miner Res 28, 2–17 (2013).

28. P. Ballanti, S. Minisola, M. T. Pacitti, L. Scarnecchia, R. Rosso, G. F. Mazzuoli, E. Bonucci, Tartrate-resistant acid phosphate activity as osteoclastic marker: sensitivity of cytochemical assessment and serum assay in comparison with standardized osteoclast histomorphometry. Osteoporos Int 7, 39–43 (1997).

29. X. Zou, J. Shen, F. Chen, K. Ting, Z. Zheng, S. Pang, J. N. Zara, J. S. Adams, C. Soo Zhang, NELL-1 binds to APR3 affecting human osteoblast proliferation and differentiation. FEBS Lett 585, 2410–2418 (2011).

30. M. Ferron, J. Wei, T. Yoshizawa, A. Del Fattore, R. A. DePinho, A. Teti, P. Ducy, G. Karsenty, Insulin signaling in osteoblasts integrates bone remodeling and energy metabolism. Cell 142, 296–308 (2010).

31. H. A. Fleisch, Bisphosphonates: preclinical aspects and use in osteoporosis. Ann Med 29, 55–62 (1997).

32. M. P. Watkins, J. Y. Norris, S. K. Grimston, X. Zhang, R. J. Phipps, F. H. Ebetino, R. Civitelli, Bisphosphonates improve trabecular bone mass and normalize cortical thickness in ovariectomized, osteoblast connexin43 deficient mice. Bone 51, 787–794 (2012).

33. K. Watanabe, A. Hishiya, Mouse models of senile osteoporosis. Mol Aspects Med 26, 221–231 (2005).

34. R. Tevlin, A. McArdle, C. K. Chan, J. Pluvinage, G. G. Walmsley, T. Wearda, O. Marecic, M. S. Hu, K. J. Paik, K. Senarath-Yapa, D. A. Atashroo, E. R. Zielins, D. C. Wan, I. L. Weissman, M. T. Longaker, Osteoclast derivation from mouse bone marrow. J Vis Exp, e52056 (2014).

35. P. Collin-Osdoby, P. Osdoby, RANKL-mediated osteoclast formation from murine RAW 264.7 cells. Methods Mol Biol 816, 187–202 (2012).

36. N. Raje, S. B. Woo, K. Hande, J. T. Yap, P. G. Richardson, S. Vallet, N. Treister, T. Hideshima, N. Sheehy, S. Chhetri, B. Connell, W. Xie, Y. T. Tai, A. Szot-Barnes, M. Tian, R. L. Schlossman, E. Weller, N. C. Munshi, A. D. Van Den Abbeele, K. C. Anderson, Clinical, radiographic, and biochemical characterization of multiple myeloma patients with osteonecrosis of the jaw. Clin Cancer Res 14, 2387–2395 (2008).

37. J. Xiang, M. A. Hurchla, F. Fontana, X. Su, S. R. Amend, A. K. Esser, G. J. Douglas, C. Mudalagiriyappa, K. E. Luker, T. Pluard, F. O. Ademuyiwa, B. Romagnoli, G. Tuffin, E. Chevalier, G. D. Luker, M. Bauer, J. Zimmermann, R. L. Aft, K. Dembowsky, K. N. Weilbaecher, CXCR4 Protein Epitope Mimetic Antagonist POL5551 Disrupts Metastasis and Enhances Chemotherapy Effect in Triple-Negative Breast Cancer. Mol Cancer Ther 14, 2473–2485 (2015).

38. J. L. Jiwoong Park, Sandeep Kumar, Damon T. Burrow, Nicholas L. Bean, Chris Chow, Nicholas C. Jacobs, Niki Song, Sarah S. Diemar, Luke A. Gilbert, Timothy R. Peterson, Using cell fitness to reduce human biases in human gene characterization. in preparation, (2019).

39. M. A. Horlbeck, A. Xu, M. Wang, N. K. Bennett, C. Y. Park, D. Bogdanoff, B. Adamson, E. D. Chow, M. Kampmann, T. R. Peterson, K. Nakamura, M. A. Fischbach, J. S. Weissman, L. A. Gilbert, Mapping the Genetic Landscape of Human Cells. Cell 174, 953–967 e922 (2018).

40. J. Bergholz, Z. X. Xiao, Role of p63 in Development, Tumorigenesis and Cancer Progression. Cancer Microenviron 5, 311–322 (2012).

41. E. L. Scheller, C. M. Baldwin, S. Kuo, N. J. D’Silva, S. E. Feinberg, P. H. Krebsbach, P. C. Edwards, Bisphosphonates inhibit expression of p63 by oral keratinocytes. J Dent Res 90, 894–899 (2011).

42. S. Han, Q. Lu, N. Wang, Apr3 accelerates the senescence of human retinal pigment epithelial cells. Mol Med Rep 13, 3121–3126 (2016).

43. E. Lecona, O. Fernandez-Capetillo, Targeting ATR in cancer. Nat Rev Cancer, (2018).

44. D. Yamada, R. Perez-Torrado, G. Filion, M. Caly, B. Jammart, V. Devignot, N. Sasai, P. Ravassard, J. Mallet, X. Sastre-Garau, M. L. Schmitz, P. A. Defossez, The human protein kinase HIPK2 phosphorylates and downregulates the methyl-binding transcription factor ZBTB4. Oncogene 28, 2535–2544 (2009).

45. A. Weber, J. Marquardt, D. Elzi, N. Forster, S. Starke, A. Glaum, D. Yamada, P. A. Defossez, J. Delrow, R. N. Eisenman, H. Christiansen, M. Eilers, Zbtb4 represses transcription of P21CIP1 and controls the cellular response to p53 activation. EMBO J 27, 1563–1574 (2008).

46. J. Y. Chou, B. C. Mansfield, The SLC37 family of sugar-phosphate/phosphate exchangers. Curr Top Membr 73, 357–382 (2014).

47. M. K. Hytonen, M. Arumilli, A. K. Lappalainen, M. Owczarek-Lipska, V. Jagannathan, S. Hundi, E. Salmela, P. Venta, E. Sarkiala, T. Jokinen, D. Gorgas, J. Kere, P. Nieminen, C. Drogemuller, H. Lohi, Molecular Characterization of Three Canine Models of Human Rare Bone Diseases: Caffey, van den Ende-Gupta, and Raine Syndromes. PLoS Genet 12, e1006037 (2016).

48. A. Guerin, L. Dupuis, R. Mendoza-Londono, in GeneReviews((R)), M. P. Adam, H. H. Ardinger, R. A. Pagon, S. E. Wallace, L. J. H. Bean, K. Stephens, A. Amemiya, Eds. (Seattle (WA), 1993).

49. X. Zou, J. Shen, F. Chen, K. Ting, Z. Zheng, S. Pang, J. N. Zara, J. S. Adams, C. Soo Zhang, NELL-1 binds to APR3 affecting human osteoblast proliferation and differentiation. FEBS Lett 585, 2410–2418 (2011).

50. J. Desai, M. E. Shannon, M. D. Johnson, D. W. Ruff, L. A. Hughes, M. K. Kerley, D. A. Carpenter, D. K. Johnson, E. M. Rinchik, C. T. Culiat, Nell1-deficient mice have reduced expression of extracellular matrix proteins causing cranial and vertebral defects. Hum Mol Genet 15, 1329–1341 (2006).

51. A. W. James, J. Shen, X. Zhang, G. Asatrian, R. Goyal, J. H. Kwak, L. Jiang, B. Bengs, C. T. Culiat, A. S. Turner, H. B. Seim Iii, B. M. Wu, K. Lyons, J. S. Adams, K. Ting, C. Soo, NELL-1 in the treatment of osteoporotic bone loss. Nat Commun 6, 7362 (2015).

52. A. Jaworski, I. Tom, R. K. Tong, H. K. Gildea, A. W. Koch, L. C. Gonzalez, M. Tessier- Lavigne, Operational redundancy in axon guidance through the multifunctional receptor Robo3 and its ligand NELL2. Science 350, 961–965 (2015).

53. J. E. Carette, C. P. Guimaraes, I. Wuethrich, V. A. Blomen, M. Varadarajan, C. Sun, G. Bell, B. Yuan, M. K. Muellner, S. M. Nijman, H. L. Ploegh, T. R. Brummelkamp, Global gene disruption in human cells to assign genes to phenotypes by deep sequencing. Nat Biotechnol 29, 542–546 (2011).

54. M. Jost, Y. Chen, L. A. Gilbert, M. A. Horlbeck, L. Krenning, G. Menchon, A. Rai, M. Y. Cho, J. J. Stern, A. E. Prota, M. Kampmann, A. Akhmanova, M. O. Steinmetz, M. E. Tanenbaum, J. S. Weissman, Combined CRISPRi/a-Based Chemical Genetic Screens Reveal that Rigosertib Is a Microtubule-Destabilizing Agent. Mol Cell 68, 210–223 e216 (2017).

55. http://weissmanlab.ucsf.edu/CRISPR/CRISPR.html.

56. T. R. Peterson, S. S. Sengupta, T. E. Harris, A. E. Carmack, S. A. Kang, E. Balderas, D. A. Guertin, K. L. Madden, A. E. Carpenter, B. N. Finck, D. M. Sabatini, mTOR complex 1 regulates lipin 1 localization to control the SREBP pathway. Cell 146, 408–420 (2011).

57. T. R. Peterson, M. Laplante, C. C. Thoreen, Y. Sancak, S. A. Kang, W. M. Kuehl, N. S. Gray, D. M. Sabatini, DEPTOR is an mTOR inhibitor frequently overexpressed in multiple myeloma cells and required for their survival. Cell 137, 873–886 (2009).

58. F. A. Ran, P. D. Hsu, J. Wright, V. Agarwala, D. A. Scott, F. Zhang, Genome engineering using the CRISPR-Cas9 system. Nat Protoc 8, 2281–2308 (2013).

59. F. P. Coxon, M. H. Helfrich, R. Van’t Hof, S. Sebti, S. H. Ralston, A. Hamilton, M. J. Rogers, Protein geranylgeranylation is required for osteoclast formation, function, and survival: inhibition by bisphosphonates and GGTI-298. J Bone Miner Res 15, 1467–1476 (2000).

60. J. C. Frith, J. Monkkonen, S. Auriola, H. Monkkonen, M. J. Rogers, The molecular mechanism of action of the antiresorptive and antiinflammatory drug clodronate: evidence for the formation in vivo of a metabolite that inhibits bone resorption and causes osteoclast and macrophage apoptosis. Arthritis and rheumatism 44, 2201–2210 (2001).

61. M. Tsubaki, M. Komai, T. Itoh, M. Imano, K. Sakamoto, H. Shimaoka, T. Takeda, N. Ogawa, K. Mashimo, D. Fujiwara, J. Mukai, K. Sakaguchi, T. Satou, S. Nishida, Nitrogen-containing bisphosphonates inhibit RANKL- and M-CSF-induced osteoclast formation through the inhibition of ERK1/2 and Akt activation. J Biomed Sci 21, 10 (2014).

62. R. Kuhn, R. M. Torres, Cre/loxP recombination system and gene targeting. Methods Mol Biol 180, 175–204 (2002).

63. https://simple.wikipedia.org/wiki/Central_dogma_of_molecular_biology.

64. F. Schwenk, U. Baron, K. Rajewsky, A cre-transgenic mouse strain for the ubiquitous deletion of loxP-flanked gene segments including deletion in germ cells. Nucleic Acids Res 23, 5080–5081 (1995).

65. C. F. Lai, S. L. Cheng, G. Mbalaviele, C. Donsante, M. Watkins, G. L. Radice, R. Civitelli, Accentuated ovariectomy-induced bone loss and altered osteogenesis in heterozygous N-cadherin null mice. J Bone Miner Res 21, 1897–1906 (2006).

66. R. K. Fuchs, R. J. Phipps, D. B. Burr, Recovery of trabecular and cortical bone turnover after discontinuation of risedronate and alendronate therapy in ovariectomized rats. J Bone Miner Res 23, 1689–1697 (2008).

67. M. R. Allen, K. Iwata, R. Phipps, D. B. Burr, Alterations in canine vertebral bone turnover, microdamage accumulation, and biomechanical properties following 1-year treatment with clinical treatment doses of risedronate or alendronate. Bone 39, 872–879 (2006).

68. M. L. Bouxsein, K. S. Myers, K. L. Shultz, L. R. Donahue, C. J. Rosen, W. G. Beamer, Ovariectomy-induced bone loss varies among inbred strains of mice. J Bone Miner Res 20, 1085–1092 (2005).

69. M. D. Willinghamm, M. D. Brodt, K. L. Lee, A. L. Stephens, J. Ye, M. J. Silva, Agerelated changes in bone structure and strength in female and male BALB/c mice. Calcif Tissue Int 86, 470–483 (2010).

70. S. K. Grimston, D. B. Goldberg, M. Watkins, M. D. Brodt, M. J. Silva, R. Civitelli, Connexin43 deficiency reduces the sensitivity of cortical bone to the effects of muscle paralysis. J Bone Miner Res 26, 2151–2160 (2011).

71. T. A. Guise, Antitumor effects of bisphosphonates: promising preclinical evidence. Cancer Treat Rev 34 Suppl 1, S19–24 (2008).

72. R. Aft, M. Naughton, K. Trinkaus, M. Watson, L. Ylagan, M. Chavez-MacGregor, J. Zhai, S. Kuo, W. Shannon, K. Diemer, V. Herrmann, J. Dietz, A. Ali, M. Ellis, P. Weiss, T. Eberlein, C. Ma, P. M. Fracasso, I. Zoberi, M. Taylor, W. Gillanders, T. Pluard, J. Mortimer, K. Weilbaecher, Effect of zoledronic acid on disseminated tumour cells in women with locally advanced breast cancer: an open label, randomised, phase 2 trial. Lancet Oncol 11, 421–428 (2010).

73. B. M. Bolstad, R. A. Irizarry, M. Astrand, T. P. Speed, A comparison of normalization methods for high density oligonucleotide array data based on variance and bias. Bioinformatics 19, 185–193 (2003).

74. C. V. Gentleman R, Huber W and Hahne F (2017).

75. G. K. Smyth, Linear models and empirical bayes methods for assessing differential expression in microarray experiments. Stat Appl Genet Mol Biol 3, Article3 (2004).

76. M. Carlson. (2016).

77. J. G. Buchan, D. M. Alvarado, G. E. Haller, C. Cruchaga, M. B. Harms, T. Zhang, M. C. Willing, D. K. Grange, A. C. Braverman, N. H. Miller, J. A. Morcuende, N. L. Tang, T. P. Lam, B. K. Ng, J. C. Cheng, M. B. Dobbs, C. A. Gurnett, Rare variants in FBN1 and FBN2 are associated with severe adolescent idiopathic scoliosis. Hum Mol Genet 23, 5271–5282 (2014).

78. C. Cruchaga, C. M. Karch, S. C. Jin, B. A. Benitez, Y. Cai, R. Guerreiro, O. Harari, J. Norton, J. Budde, S. Bertelsen, A. T. Jeng, B. Cooper, T. Skorupa, D. Carrell, D. Levitch, S. Hsu, J. Choi, M. Ryten, J. Hardy, D. Trabzuni, M. E. Weale, A. Ramasamy, C. Smith, C. Sassi, J. Bras, J. R. Gibbs, D. G. Hernandez, M. K. Lupton, J. Powell, P. Forabosco, P. G. Ridge, C. D. Corcoran, J. T. Tschanz, M. C. Norton, R. G. Munger, C. Schmutz, M. Leary, F. Y. Demirci, M. N. Bamne, X. Wang, O. L. Lopez, M. Ganguli, C. Medway, J. Turton, J. Lord, A. Braae, I. Barber, K. Brown, P. Passmore, D. Craig, J. Johnston, B. McGuinness, S. Todd, R. Heun, H. Kolsch, P. G. Kehoe, N. M. Hooper, E. R. Vardy, D. M. Mann, S. Pickering-Brown, N. Kalsheker, J. Lowe, K. Morgan, A. David Smith, G. Wilcock, D. Warden, C. Holmes, P. Pastor, O. Lorenzo-Betancor, Z. Brkanac, E. Scott, E. Topol, E. Rogaeva, A. B. Singleton, M. I. Kamboh, P. St George-Hyslop, N. Cairns, J. C. Morris, J. S. Kauwe, A. M. Goate, Rare coding variants in the phospholipase D3 gene confer risk for Alzheimer’s disease. Nature 505, 550–554 (2014).

79. P. Krawitz, C. Rodelsperger, M. Jager, L. Jostins, S. Bauer, P. N. Robinson, Microindel detection in short-read sequence data. Bioinformatics 26, 722–729 (2010).

80. A. McKenna, M. Hanna, E. Banks, A. Sivachenko, K. Cibulskis, A. Kernytsky, K. Garimella, D. Altshuler, S. Gabriel, M. Daly, M. A. DePristo, The Genome Analysis Toolkit: a MapReduce framework for analyzing next-generation DNA sequencing data. Genome Res 20, 1297–1303 (2010).

81. G. A. Van der Auwera, M. O. Carneiro, C. Hartl, R. Poplin, G. Del Angel, A. Levy- Moonshine, T. Jordan, K. Shakir, D. Roazen, J. Thibault, E. Banks, K. V. Garimella, D. Altshuler, S. Gabriel, M. A. DePristo, From FastQ data to high confidence variant calls: the Genome Analysis Toolkit best practices pipeline. Curr Protoc Bioinformatics 43, 11 10 11–33 (2013).

82. M. Lek, K. J. Karczewski, E. V. Minikel, K. E. Samocha, E. Banks, T. Fennell, A. H. O’Donnell-Luria, J. S. Ware, A. J. Hill, B. B. Cummings, T. Tukiainen, D. P. Birnbaum, J. A. Kosmicki, L. E. Duncan, K. Estrada, F. Zhao, J. Zou, E. Pierce-Hoffman, J. Berghout, D. N. Cooper, N. Deflaux, M. DePristo, R. Do, J. Flannick, M. Fromer, L. Gauthier, J. Goldstein, N. Gupta, D. Howrigan, A. Kiezun, M. I. Kurki, A. L. Moonshine, P. Natarajan, L. Orozco, G. M. Peloso, R. Poplin, M. A. Rivas, V. Ruano-Rubio, S. A. Rose, D. M. Ruderfer, K. Shakir, P. D. Stenson, C. Stevens, B. P. Thomas, G. Tiao, M. T. Tusie-Luna, B. Weisburd, H. H. Won, D. Yu, D. M. Altshuler, D. Ardissino, M. Boehnke, J. Danesh, S. Donnelly, R. Elosua, J. C. Florez, S. B. Gabriel, G. Getz, S. J. Glatt, C. M. Hultman, S. Kathiresan, M. Laakso, S. McCarroll, M. I. McCarthy, D. McGovern, R. McPherson, B. M. Neale, A. Palotie, S. M. Purcell, D. Saleheen, J. M. Scharf, P. Sklar, P. F. Sullivan, J. Tuomilehto, M. T. Tsuang, H. C. Watkins, J. G. Wilson, M. J. Daly, D. G. MacArthur, C. Exome Aggregation, Analysis of protein-coding genetic variation in 60,706 humans. Nature 536, 285–291 (2016).

